# Whole genome comparisons of *Staphylococcus agnetis* isolates from cattle and chickens

**DOI:** 10.1101/2020.01.06.896779

**Authors:** Abdulkarim Shwani, Pamela R. F. Adkins, Nnamdi S. Ekesi, Adnan Alrubaye, Michael J. Calcutt, John R. Middleton, Douglas D. Rhoads

## Abstract

*S. agnetis* has been previously associated with subclinical or clinically mild cases of mastitis in dairy cattle and is one of several Staphylococcal species that have been isolated from the bone and blood of lame broilers. We were the first to report that *S. agnetis* could be obtained frequently from bacterial chondronecrosis with osteomyelitis (BCO) lesions of lame broilers. Further, we showed that a particular isolate of *S. agnetis*, chicken isolate 908, can induce lameness in over 50% of exposed chickens, far exceeding normal BCO incidences in broiler operations. We have previously reported the assembly and annotation of the genome of isolate 908. To better understand the relationship between dairy cattle and broiler isolates, we assembled 11 additional genomes for *S. agnetis* isolates, including an additional chicken BCO strain, and ten isolates from milk, mammary gland secretions or udder skin, from the collection at the University of Missouri. To trace phylogenetic relationships, we constructed phylogenetic trees based on multi-locus sequence typing, and Genome-to-Genome Distance Comparisons. Chicken isolate 908 clustered with two of the cattle isolates along with three isolates from chickens in Denmark and an isolate of *S. agnetis* we isolated from a BCO lesion on a commercial broiler farm in Arkansas. We used a number of BLAST tools to compare the chicken isolates to those from cattle and identified 98 coding sequences distinguishing isolate 908 from the cattle isolates. None of the identified genes explain the differences in host or tissue tropism. These analyses are critical to understanding how Staphylococci colonize and infect different hosts and potentially how they can transition to alternative niches (bone vs dermis).

**Importance:** *Staphylococcus agnetis* has been recently recognized as associated with disease in dairy cattle and meat type chickens. The infections appear to be limited in cattle and systemic in broilers. This report details the molecular relationships between cattle and chicken isolates in order to understand how this recently recognized species infects different hosts with different disease manifestations. The data show the chicken and cattle isolates are very closely related but the chicken isolates all cluster together suggesting a single jump from cattle to chickens.

## Introduction

In the US, skeletal problems are estimated to cost the broiler industry more than 100 million dollars annually (1-5). Lameness is an important chicken industry issue affecting from 1-10% of a flock, and a wide array of bacterial genera have been isolated from chickens affected by bacterial chondronecrosis with osteomyelitis (BCO) (5-25). *Staphylococcus agnetis*, a coagulase-variable, Gram-positive bacterium has been found to cause infections of the bones and blood of broilers leading to BCO (26, 27). BCO primarily affects the growth plate in the proximal femur and tibia, the fast-growing leg bones. We have shown that an isolate of *S. agnetis* (strain 908) obtained from BCO chickens can induce BCO lameness at levels greater than 50% of the population when administered in a single dose in drinking water (26, 27). Previously, *S. agnetis* has also been associated with subclinical or mild cases of clinical mastitis in dairy cattle (28-31). There are very few reports of *S. agnetis* in poultry and we have speculated that the virulent strain we isolated may be the result of prolonged selection resulting from years of inducing BCO lameness at our research farm. Genome sequence analysis of multiple isolates of *S. agnetis* from the University of Arkansas research farm have revealed little sequence variation and thus they appear to be clonal (unpublished data). The annotated complete genome of strain 908 has been published (26). Draft genomes of a cattle isolate, *S. agnetis* CBMRN20813338 (32), and chicken isolates (33) have been deposited in the NCBI genome databases. To better understand the phylogenomic relationships between dairy cattle and broiler isolates, we have generated genome assemblies for multiple cattle isolates and an additional chicken isolate of *S. agnetis*. We used multi-locus sequence typing (MLST) and genome distance comparisons to develop phylogenetic trees. We also performed reciprocal BLAST and BLASTX comparisons to identify genes and gene islands that distinguish the chicken and cattle isolates. The goal was to determine the phylogenetic relationships between cattle and chicken isolates, and whether there were easily discernable genes responsible for the virulence of isolate 908, or species-specific pathogenesis.

## Results

### *Staphylococcus agnetis* genomes assemblies

Sources (host, tissue, disease) for the *S. agnetis* isolates used in our analyses are presented in Table 1. For this work, we generated draft genomes for eight cattle isolates (1383, 1384, 1385, 1387, 1389, 1390, 1391, 1392) from 2×251 paired end MiSeq reads (Table S1), and we generated one finished genome for one cattle isolate (1379). These cattle isolates were cultured from skin swabs (1385, 1389), milk (1379, 1383, 1384, 1387, 1391, 1392), or pre-partum mammary gland secretions (1390). The new, draft, *de novo* assemblies for the eight cattle isolates ranged from 43 to 328 contigs comprising 2.381 to 2.581 Mbp (Table S1). The hybrid assembly from long and short reads (see Materials and Methods for details) for cattle isolate 1379 produced a single chromosome of 2.45 Mbp. We identified *S. agnetis* isolate 1416 from a BCO lesion in a necropsy sampling of BCO birds on a commercial broiler farm in Arkansas. The hybrid assembly of the 1416 genome produced a 2.45 Mbp chromosome and plasmids of 59 and 28 kbp. We included the draft genome for cattle isolate CBMRN20813338 as it was the first *S. agnetis* genome characterized (32). We had earlier published the finished genome for chicken isolate *S. agnetis* 908, a *de novo* assembly of Pacific Biosciences long reads, with subsequent correction with MiSeq reads (26). This assembly includes a single 2.47 Mbp chromosome and a 29 kbp plasmid. Recently, we identified two additional plasmids of 3.0 and 2.2 kbp (unpublished) from the assembly data that we have included in our genome comparisons.

**Table 1.**
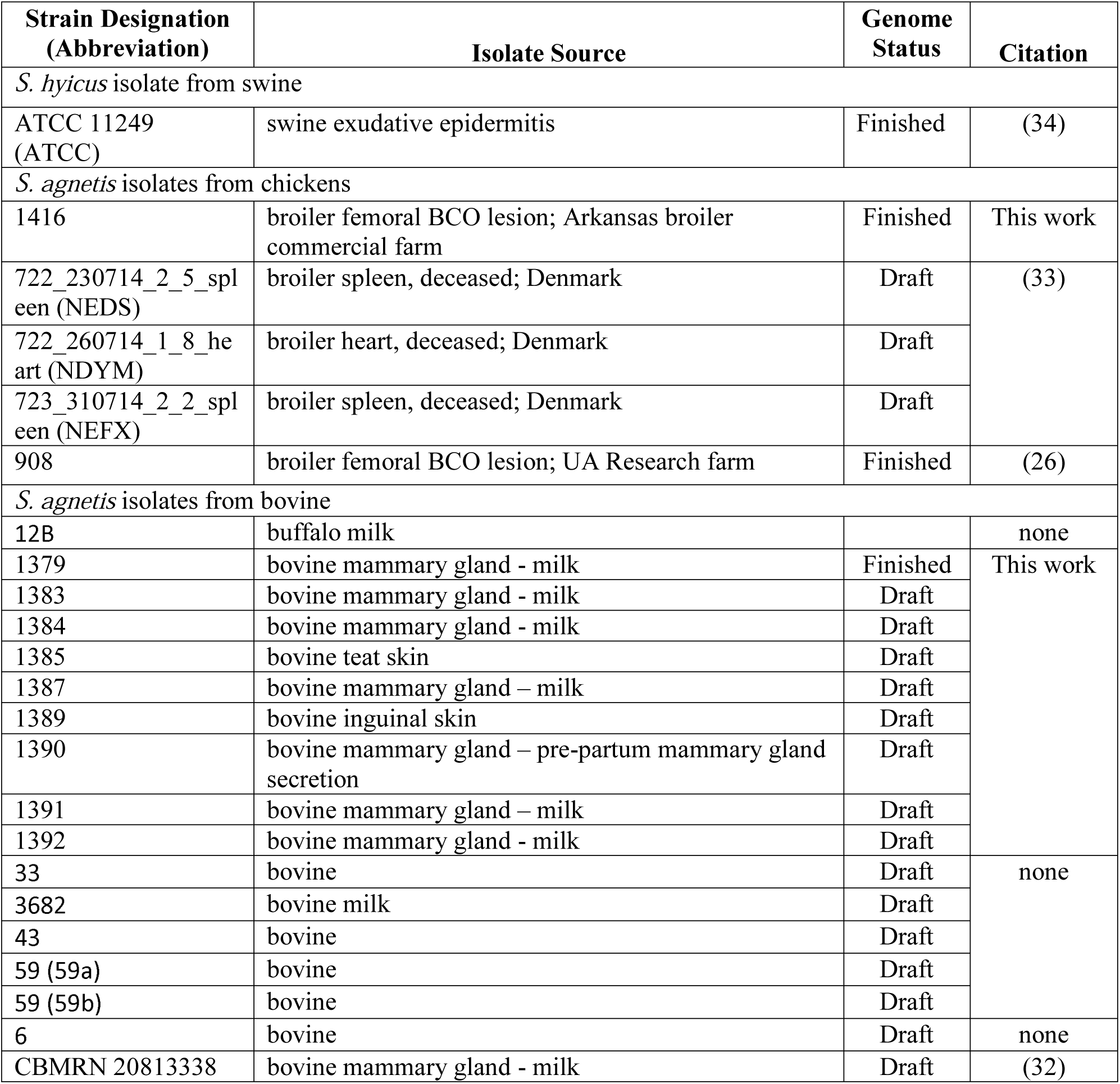
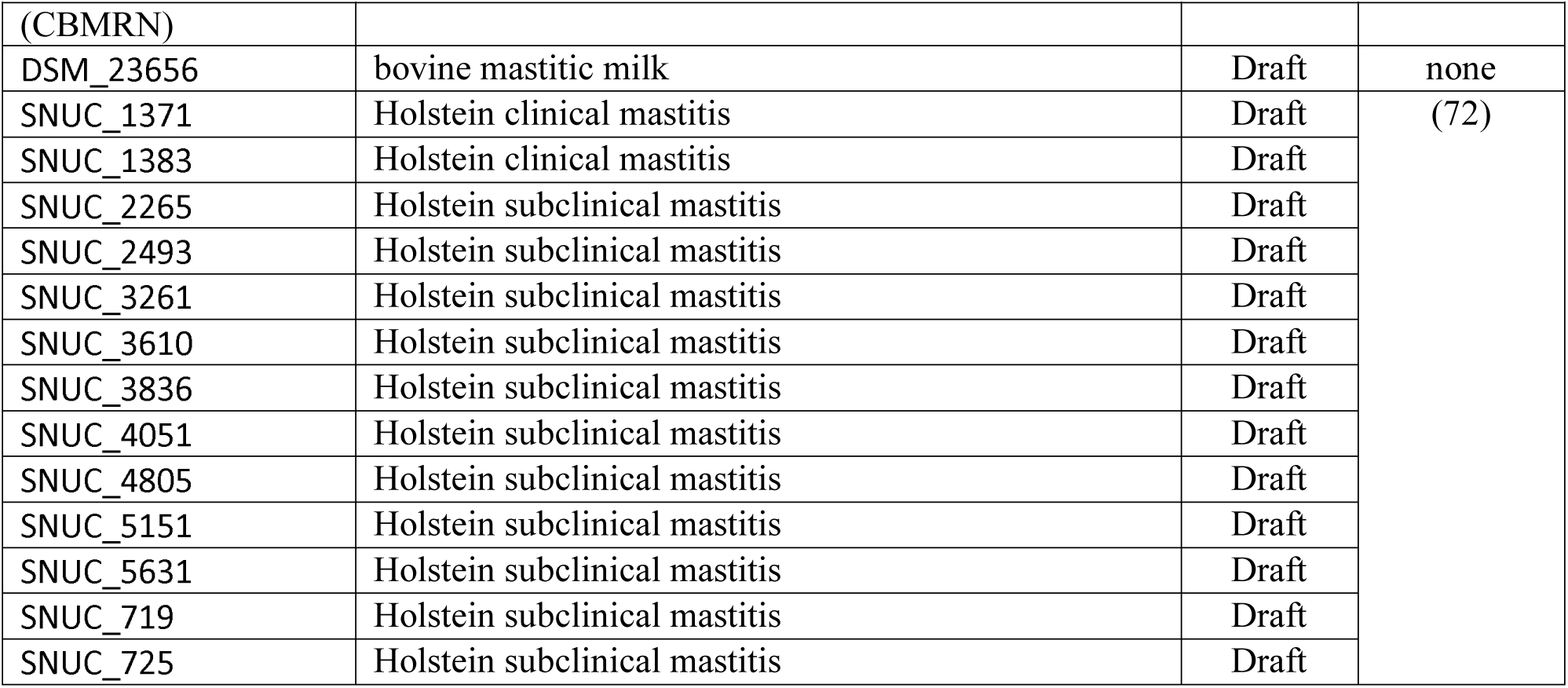
Bacterial genomes utilized in these analyses. Genomes are separated by species designation and host source. For each genome the isolate designation is given as well as our abbreviation for this manuscript where indicated. The Isolate Source indicates tissue or sample source for the bacterial isolate. Genome status indicates whether the genome is considered finished or draft. Citation is the publication source. Further information on the genome assemblies is provided in Table S1.

### Phylogenetic analyses

To begin to trace the phylogenomic relationship between the cattle *S. agnetis* isolates and those from chicken, we first generated MLST phylogenetic trees. We included a total of 13 isolates, including the published genome for cattle isolate *S. agnetis* CBMRN20813338, the 9 new cattle isolate assemblies, chicken isolates 908 and 1416, and *S. agnetis* isolate 12B from NCBI. We included the genome from strain 12B, isolated from the milk of a buffalo with bubaline mastitis. The dendrogram from genome BLAST on the NCBI genome page for *S. agnetis*, presents 12B as the closest genome for a bovine isolate to the genome for chicken isolate 908. *S. hyicus* ATCC11249^T^ (34) from swine exudative dermatitis was used as the outgroup. Figure 1 presents a tree based on seven housekeeping genes (*ackA, fdhD, fdhF, grol, purA, tpiA, tuf*), where orthologs could be identified in each of the assemblies, and the genes are dispersed throughout the 908 chromosome. From the MLST phylogenetic tree, we see two chicken isolates (908 and 1416) cluster within the cattle *S. agnetis* isolates within a clade with bovine isolates 1379, 1387 and 12B. MLST analysis with seven virulence genes (encoding five distinct fibronectin binding proteins and two exotoxins) identified in all of the assemblies produced a tree with a very similar topology (data not shown).

**Figure 1.**
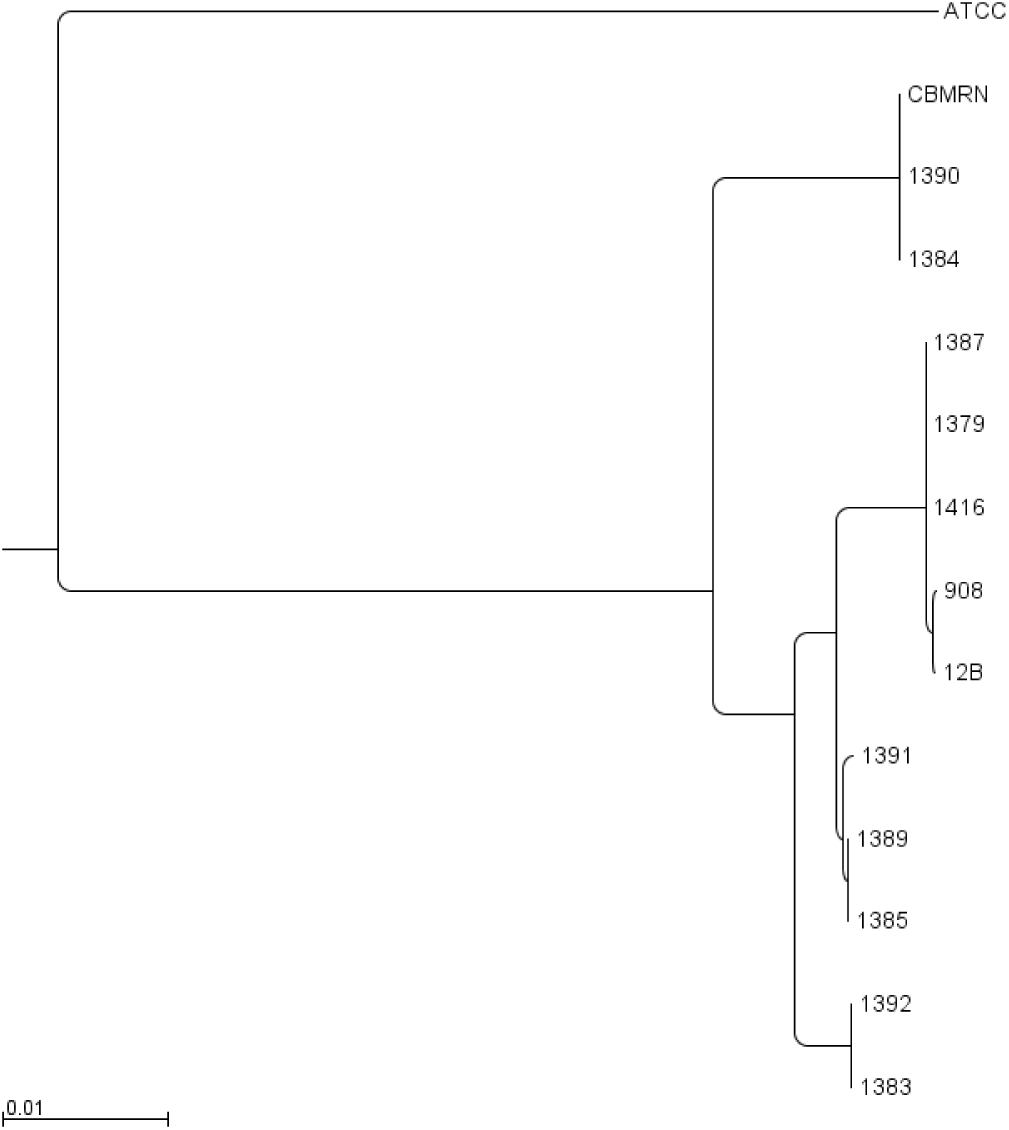
MLST phylogeny two chicken and eleven bovine, isolates of *S. agnetis*, with a swine *S. hyicus* (ATCC) isolate as the out-group. The tree is based on seven housekeeping genes (see text).

Since we began our analyses, additional genomes for isolates of *S. agnetis* have been deposited in NCBI. The NCBI dendrogram based on genomic BLASTN (https://www.ncbi.nlm.nih.gov/genome/?term=agnetis) for 26 *S. agnetis* assemblies, indicates that our 908 chicken isolate clusters with three Danish chicken isolates and 5 bovine isolates (12B, SUNC_2265, SNUC_4805, SNUC_5151, SUC_3261) in a single clade relative to CBMRN20813338, and 16 additional bovine isolates. In order to expand on the MLST analyses and all 36 genomes (26 in NCBI and our 10 new assemblies, including 9 isolates of bovine origin and 1 isolate of chicken origin) we used the Genome-to-Genome Distance Calculator (GGDC) to generate a phylogenetic tree based on genetic distances computed from whole genome BLASTN comparisons (35). This included our two chicken isolates (908, 1416) and three chicken isolates (NEDS, NEFX, NDYM) from organs from two deceased broilers on a farm in Denmark (33). The phylogenetic tree based on genome distances (Fig. 2) shows that four of the genomes from chicken isolates (908, NEDS, NEFX, NDYM) cluster within the cattle isolates with the Denmark chicken isolates being most similar to our chicken isolate 908. Chicken isolate 1416 is in a sister branch clustered with 7 bovine isolates including isolate 12B from the milk of a buffalo in Argentina with mastitis. The data are consistent with five, or potentially six, different clades within the *S. agnetis* species group with the five chicken isolate genomes all within one clade. The nine new genomes for mastitis-related isolates of *S. agnetis* from the USA are distributed across all branches of the tree. There is no indication of geographic restriction of particular genotypes for *S. agnetis* isolated from the bovine mammary gland. Nor is there a particularly noticeable separation of the chicken isolate genomes from the cattle isolate genomes. We also analyzed all of the genomes by Average Nucleotide Identity (36) and obtained the same phylogenomic architecture (data not shown).

**Figure 2.**
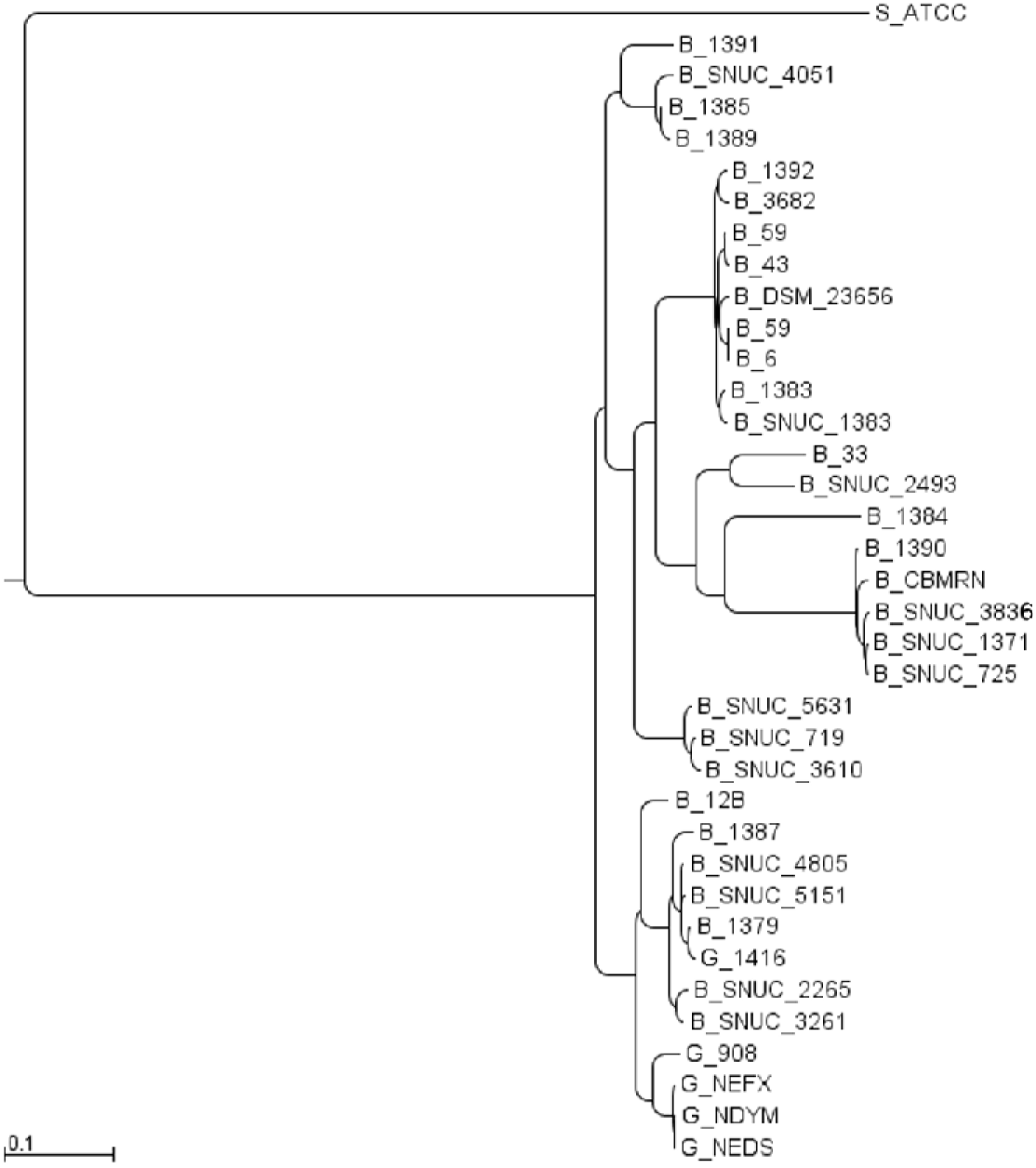
Genome to Genome Distance phylogenetic tree for 35 *S*.*agnetis* genomes. GGDC data (formula 2) was used to construct a neighbor-joining tree for comparison of five chicken (G_) and 31 bovine (B_) isolates, of *S. agnetis*. Isolates are described in the text and Table S1. *S. hyicus* (ATCC) from swine (S_) was the out group.

### Genome Comparison

We used CGView Server (37) to perform and visualize comparisons of the 2.47 Mbp chromosome from chicken isolate 908 to three cattle isolate genomes; the finished 1379 isolate genome and draft genomes for isolates 1387 and 1385 (Fig. 3). We selected the 1379 isolate genome for production of a finished genome based on being one of the closest genomes to the chicken isolates (Fig. 2). We included the draft 1387 isolate genome from the same branch as the chicken isolates, and 1385 as the largest assembly of the other cattle isolates. The CGView in Figure 3 identified five gene islands which appear to distinguish chicken isolate 908, from the three cattle isolates. The five islands were also visible when we compared chicken isolate 908 to our other new draft cattle assemblies or the buffalo isolate 12B (data not shown). We hypothesized that these islands could potentially contain sequences related to host adaptation. We annotated the genes in these five islands using BLASTP and further evaluated for presence in the other four chicken isolate genomes, or any of the currently available 36 bovine isolate genomes (Table S2). Regions in the 908 genome not represented in the cattle isolates according to the CGView are located approximately as follows: island 1 for 167-235 kbp; island 2 for 978-1021 kbp; island 3 for 1162-1177 kbp; island 4 for 1831-1848 kbp; and island 5 for 2007-2018 kbp. Thus, approximately 154 kbp out of 2474 kbp are distinct from the cattle isolates.

**Figure 3.**
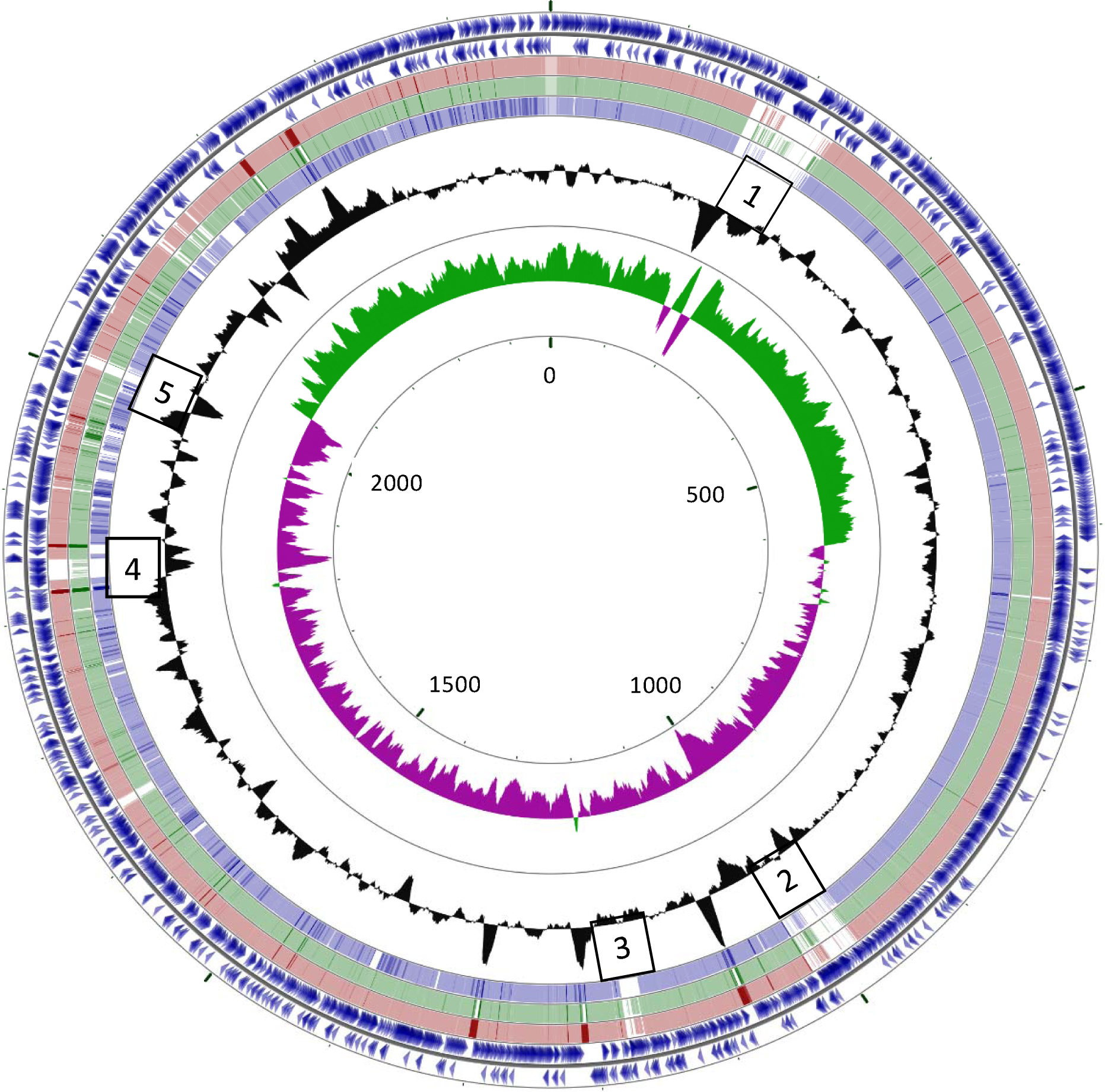
CGView blastn comparison of *S. agnetis* 908, 1379, 1387 and 1385 genomes. The *S. agnetis* 908 2.4 Mbp chromosome was the reference for blastn comparisons with three *S. agnetis* cattle isolates, 1379 (pink), 1387 (green), and 1385 (blue). Parameters were Query size = 10000, overlap 5000, expect=0.0001. Intensity of the color is indicative of the blastn score. The outer two rings show the annotated genes for isolate *S. agnetis* 908. The innermost ring indicates Mbp, the second most innner ring is the GC skew (magenta GC skew-; green GC skew+), and the third most inner ring plots GC content. The Numbered boxes indicate the locations of the 5 gene islands discussed in the text and are not scaled to the size of the island.

Analysis of the 908 2.47 Mbp chromosome for prophage using the PHASTER website (data not shown) identifies island 1 as containing two intact Staphylococcal prophage (Staphy_EW_NC_007056 and Staphy_IME_SA4_NC_029025) from 170.1 to 232.7 kbp. Islands 2 and 3 are identified as questionable for being complete prophage. Island 2 is Staphy_2638A_NC_007051 from 980.8 to 1020.9 kbp and island 3 is most similar to a *Clostridium* phage, phiMMP04_NC_019422, from 1156.8 to 1178.5 kbp. Island 4 contains genes indicative of a conjugative transposon, with sequence similarity to Tn*6012* of *S. aureus*, inserted in an intergenic region approximately 170 bp upstream of the *bioD* gene. The shortest of the five blocks is island 5 (∼11 kbp), which contains an apparent operon encoding a strain variable Type 1 DNA restriction-modification system (*hsdMSR*). Therefore, 124.4 of the estimated 154 kbp in the five islands represents prophage sequences, while the other two contain a probable transposon and a restriction-modification operon. The most similar match to island 4 and 5 in BLAST searches at NCBI were to *S. aureus* genomes. Figure 4 further relates these 5 islands as candidates for host adaptation by comparing the 908 isolate 2.47 Mbp chromosome to the finished 1379 bovine isolate genome and draft assemblies of chicken isolates NEDS from the a deceased broiler in Denmark and the finished genome assembly of isolate 1416 from a BCO broiler on a commercial farm in Arkansas. From these comparisons we conclude that islands 1, 2, 4, and 5 are, for the most part, present in at least one of the other two chicken isolates, but none of the islands is in both of the other chicken isolates (i.e., specific to all chicken isolates). There is the caveat that the 908 isolate and 1416 isolate genomes are finished genomes and the NEDS genome is only a draft genome so any island not found in the NEDS assembly could potentially be an assembly issue. Close inspection of the TBLASTN analysis of the proteins from the islands for presence in any of the bovine isolate genomes (Table S2) 164 of the 908 predicted polypeptides have significant matches in at least one of the cattle isolates, while only 32 are not found in any of the cattle isolates. For those 32 polypeptides, 31 are also not found in either of the four other chicken isolates (1416 or the three Danish isolates). Polypeptide 217 is the only polypeptide not identified in any bovine isolate genome but is identified in the 1416 genome. Polypeptide 217 is a 47 amino acid hypothetical protein, so we see no real islands of polypeptides (i.e., chicken specific pathogenicity island) that distinguish chicken isolate genomes from the bovine isolate genomes.

**Figure 4.**
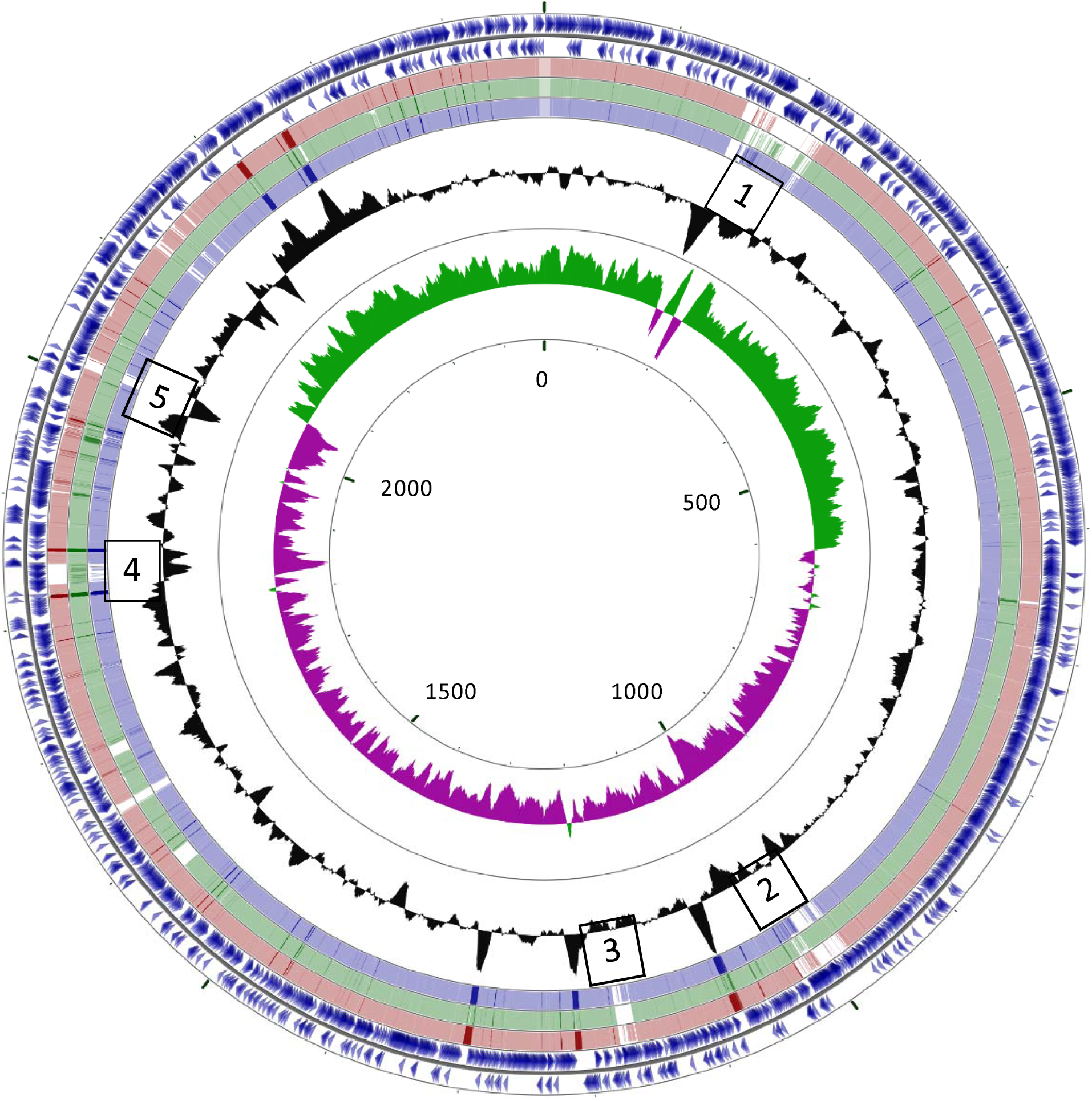
CGView blast comparison of *S. agnetis* 908, 1379, NEDS, and 1416 genomes. The *S. agnetis* 908 2.4 Mbp chromosome was the reference for blastn comparisons with *S. agnetis* cattle isolate 1379 (pink), and chicken isolates; NEDS (green), and 1416 (blue). All other details are as in Figure 3 legend.

We have assembled sequences of three plasmids in isolate 908 (29, 3 and 2.2 kbp). None of these plasmids is found in any of our nine newly assembled cattle isolate listed in Table 1. There are presently 26 genome assemblies for *S. agnetis* in the NCBI database. Only two assemblies are listed as “completed” (i.e., finished), our assembly for chicken isolate 908 (26), and isolate 12B from buffalo milk in Argentina (unpublished). The other 24 are draft assemblies. The NCBI genomes include 21 isolates from cattle, one from buffalo, and four chicken isolates; 908 and the three chicken isolates from Denmark (NEDS, NEFX, and NDYM). We performed a BLASTN search using the NCBI program selection “optimize for highly similar sequences (megablast)” where the query was the three plasmids from isolate 908 plasmids and the database searched was the 26 assemblies already in NCBI. The 29 kbp plasmid identified one 4729 base contig in the NEFX assembly with 22% query coverage in 10 different regions, with the largest region comprising 3317 identities over 3331 bases. The 3 kbp plasmid matched 450 out of 478 bases in 565 bp contigs in all three of the Danish chicken isolate assemblies (NDYM, NEDS, NEFX). The 2.2 kbp plasmid matched 2071 out of 2080 bases in a 2304 base contig in the chicken NEFX assembly. We also performed BLASTN searches of the three plasmids using the NCBI NR database exclusive of *S. agnetis* entries. The best matches for the 29 kbp plasmid are to the 30.9 kbp plasmid pH1-1 from a pheasant isolate of *S. aureus*. The two plasmids share 99% identity with 39% query coverage in three different regions of the plasmid (2520, 1573, and 982 bp). The 3 kbp 908 plasmid has a 43% query cover with 89% identity with an unnamed 37.2 kbp *S. aureus* plasmid from a human isolate of *S. aureus*. The best match for the 2.2 kbp 908 plasmid was 99% identity for 2076 bp in a 46.5 kbp plasmid pSALNT46 from an *S. aureus* isolate from retail turkey meat. We therefore conclude that none of the plasmids in isolate 908 appear to correspond to a chicken host specialization determinant for the jump from to chickens, but some sequences in the 29 kbp plasmid and the 2.2 kbp plasmid could have been picked up after the jump to poultry.

To screen at higher resolution, we used the SEED Viewer sequence comparison tool to compare entire assemblies for individual polypeptide coding sequences for four isolates: chicken isolate 908 (including the 3 plasmids), chicken isolate 1416, bovine isolate 1379 and the bovine CBMRN isolate. We used isolates 908 and 1416 as finished assemblies of two chicken isolates, 1379 as a finished assembly of a cattle isolate, and CBMRN as the original draft cattle isolate. The BLASTP comparison results were then filtered for isolate 908 polypeptides with >90% identity for polypeptides in isolate 1416, but <50% identity for isolate 1379 and the CBMRN isolate. This filter identified 99 polypeptides (Table S3). To predict functions of these 99 polypeptides we used both the RAST annotation and the NCBI Prokaryote Genome Annotation Pipeline (PGAP) to categorize the potential function of these 99 polypeptides. We identified 75 polypeptides as hypothetical or of unknown function, 11 phage related, and 8 involved in mobile elements or plasmid maintenance. The remaining five polypeptides, listed under “other”, are: deoxyuridine 5’-triphosphate nucleotidohydrolase (EC 3.6.1.23), hypothetical SAR0365 homolog in superantigen-encoding pathogenicity islands SaPI, ribosyl nicotinamide transporter PnuC-like, aspartate aminotransferase (EC 2.6.1.1), and N-acetyl-L,L-diaminopimelate aminotransferase (EC 2.6.1.-). If we relaxed the cutoff for the cattle isolates to <70% identity in cattle isolates we identified 9 additional polypeptides which added one additional hypothetical polypeptide, four additional phage related polypeptides, and four additional polypeptides involved in mobile element or plasmid maintenance. There were no additional polypeptides in the “other” category, only the five described above (Table S3).

The dUTP nucleotidohydrolase (Gene ID 209; 191,107-191,625 bp) is annotated as a phage related protein with roles in viral replication for reducing incorporation of uracil in viral DNA and is located within the Staphy_EW_NC_007056 prophage in island 1 described above, so this gene is likely to function primarily in the biology of that prophage.

The SAR0365 homolog (Gene ID 1037; 1,018,987-1,020,928 bp) is encoded in island 2 within the Staphy_2638A_NC_007051 prophage. SAR0365 polypeptide is a hypothetical protein that the NCBI Prokaryotic Genome Annotation Pipeline annotates as a toxin in the PemK/MazF type II toxin-antitoxin system. Four of the seven protein entries for SAR0365 homologs in NCBI are associated with superantigen-encoding pathogenicity islands (SaPI) in clinical isolates of *Staphylococcus aureus* and two are associated with *S. aureus* phages. Mobilization of SaPI has been associated with temperate phage replication (38). The 908 isolate genome contains additional hypothetical genes annotated as SaPI-associated homologs (Gene ID 1184 1,173,209-1,173,412 bp; Gene ID 1185 1,173,409-1,173,714 bp; Gene ID 1938 1,957,610-1,959,703 bp; Gene ID 1939 1,959,935-1,961,452 bp; Gene ID 2116 2,136,487-2,136,603 bp). Mobilization depends on a terminase (38), but the only SaPI associated terminase is Gene ID 2092 (2,118,053-2,118,355 bp). We had previously described a cluster of five exotoxin/superantigen-like proteins from 1,956,884 to 1,968,958 bp (26). Therefore, the only potential superantigen-containing pathogenicity island would approximate from 1.95 to 2.12 Mbp which would be larger than the prototypical 15-18 kbp SaPI (38). A BLASTP of the *S. agnetis* protein database at NCBI with the predicted protein for Gene ID 1037 (SAR0365 homolog) identified the isolate 908 entry, as well as identical entries in all three Danish chicken isolates (NEDS, NDYM, NEFX), but no entries in any of the 20 cattle *S. agnetis* isolates in NCBI. Expanding the BLASTP to all Staphylococcaceae identified highly similar matches (92% identity, 100% query coverage) in *Staphylococcus hominis* and less similar (50% identity, 98% query coverage) in *S. aureus*. Further analyses and additional samples would be required to speculate further regarding the role of this SAR0365 homolog as a virulence factor in chicken tropism.

The genes for ribosyl nicotinamide transporter (Gene ID 2466), aspartate aminotransferase (Gene ID 2469) and N-acetyl-L,L-diaminopimelate aminotransferase (Gene ID 2470) are located in a five gene region on the 29 kbp plasmid, with the other 2 genes encoding hypothetical polypeptides. We performed a BLASTP search of the Staphylococcaceae proteins in the NCBI database. The 89 amino acid ribosyl nicotinamide transporter matched multiple entries from *S. aureus* and all three Danish *S. agnetis* isolates from chicken. Many of the BLASTP hits for this polypeptide are annotated as an AbrB family transcriptional regulator by the NCBI Prokaryotic Genome Annotation Pipeline (PGAP). The 64 amino acid hypothetical polypeptide for Gene ID 2467 is only conserved in two entries from the Danish *S. agnetis* broiler isolates. The 197 residue polypeptide from Gene ID 2468 is well conserved in a broad swath of staphylococci and PGAP annotates this polypeptide as an IS*6* family transposase. We note that Gene ID 2469 and 2470 are close to each other and in different reading frames suggestive of a possible frameshift introduced as an assembly error. Indeed, if we join the predicted polypeptides of these two ORFs, BLASTP analysis identifies *Staphylooccus* protein entries that match over the span of the merged polypeptides. However, we have evaluated this hypothesis further by templated assembly of the 908 MiSeq data onto the 29 kbp plasmid sequence. The MiSeq reads all agree with the assembly as presented in our NCBI submission. Therefore, these two ORFs in the 29 kbp plasmid may be a frameshifted pseudogene or, if translated may have alternate functions for this organism.

Finally, to determine whether there were any regions in the cattle isolates that are not found in the chicken isolate genome assemblies, we performed a CGView analysis with the finished genome from isolate 1379 as the reference (Fig. 5). We included cattle isolate CBMRN and compared to the finished genomes of chicken isolates 908 (chromosome plus three plasmids), and 1416. There were cattle isolate regions that appeared to be absent from one of the chicken isolates but there were no regions found in both cattle isolates that were missing from both chicken isolates.

**Figure 5.**
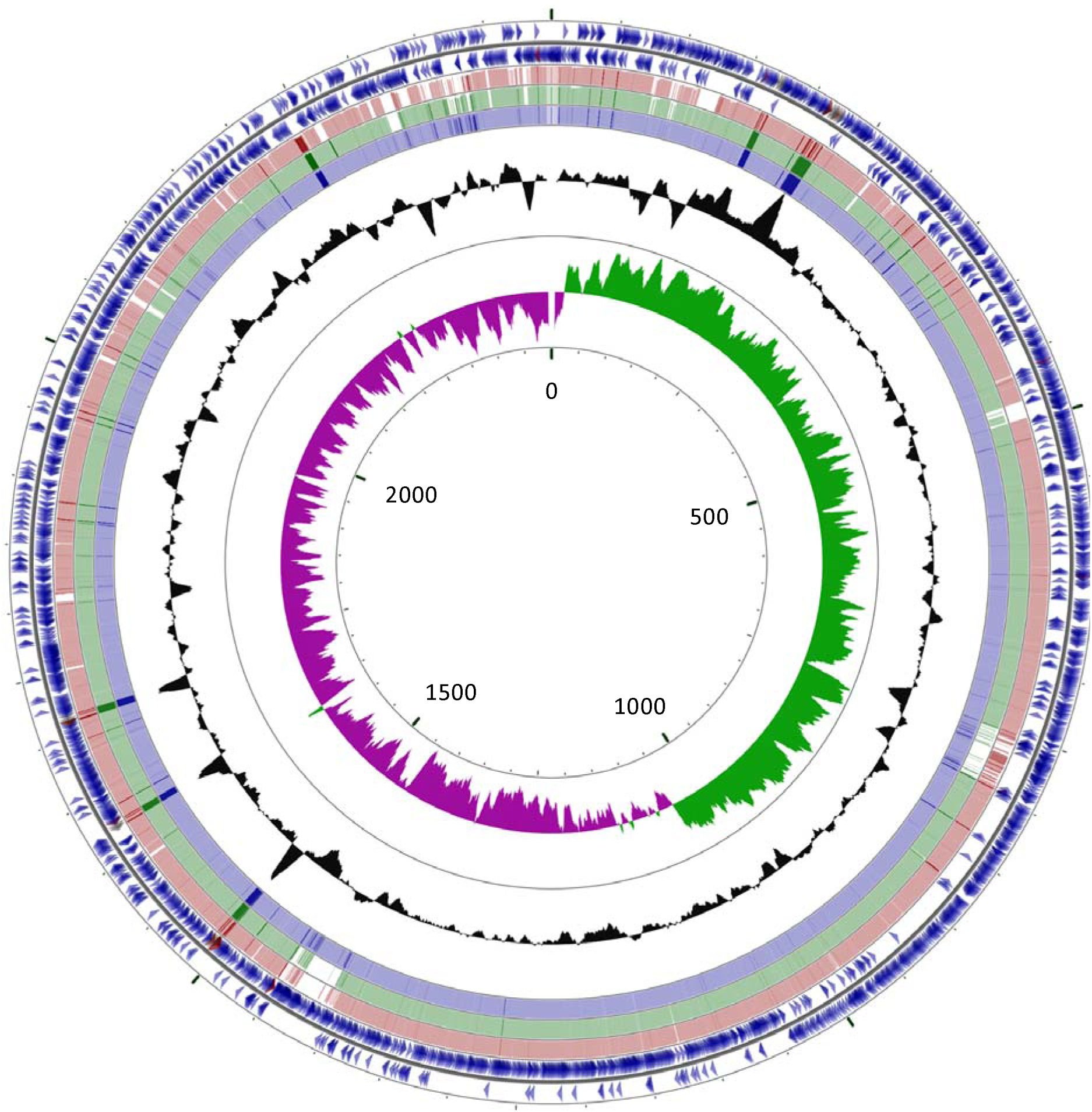
CGView blastn comparison of *S. agnetis* 1379, CBMRN, 908 and 1416 genomes. The *S. agnetis* 1379 2.4 Mbp chromosome was the reference for blastn comparisons with *S. agnetis* cattle isolate CBMRN (pink), and chicken isolates; 908 (green), and 1416 (blue). All other details are as in Figure 3 legend.

## Discussion

The Staphylococcal genus not only includes a number of pathogenic species infecting vertebrate animals worldwide, but also includes many saprophytic or commensal species (39-41). *S. agnetis* is closely related to *S. hyicus* and *S. chromogenes* and was only described as a distinct staphylococcal species in 2012, based on DNA sequence differences of rDNA and two protein coding genes in isolates from mastitis in dairy cattle (42). *S. agnetis* cannot be easily differentiated from *S. hyicus* using routine speciation techniques, e.g. partial 16S rDNA sequencing, MALDI-TOF, or fermentation methods (28-30, 43). Hence, *S. agnetis* has either escaped recognition due to misclassification or is an emerging pathogen in some agricultural animal species. While *S. agnetis* was originally reported in cattle mastitis (42), it has more recently been reported in chicken bone infections and in multiple internal organs from deceased broilers (33). Metagenomics also detected *S. agnetis* 16S rDNA sequences in the gut of a sheep scab mite (44). Phylogenetic analyses based on 16S rRNA sequences cluster *S. agnetis* very close to *S. hyicus* (26) with a group of staphylococci associated with domestic vertebrate species (e.g., cattle, swine, dog) (28-31, 34, 45-53). Most of these species are associated with dermal or epithelial infections, such as exudative dermatitis (28, 29, 31, 34, 47, 49-51, 53-55) and not with osteomyelitis as we have seen with chicken isolate 908 (26, 27). The more phylogenetically distant taxon, *S. aureus*, is prominently known for osteomyelitis in humans (56-58). The Danish broiler chicken isolates were from multiple tissues from deceased birds and we have no information about possible osteomyelitis. That the three Danish *S. agnetis* isolates, and our isolates 908 and 1416, are all closely related and within a clade of the cattle *S. agnetis* isolates suggests a recent expansion of the host range (i.e. from cattle to chickens) as seen for a human-specific clade of *S. aureus* that “jumped” to chickens in the United Kingdom (17). A single radiation out of the cattle group also argues against *S. agnetis* jumping back and forth between cattle and chickens. We have previously reported that isolate 908 can produce a bacteremia in the latter stages of BCO development before the birds are overtly lame (5, 26, 27, 59). We do not know if the Danish isolates can induce the BCO lameness that we have demonstrated for isolate 908 (26, 27). Our isolate 908 appears to represent a hypervirulent clone expanded through years of inducing high levels of BCO lameness on our research farm (5, 26, 27, 60-63) and could have evolved through selection from less virulent *S. agnetis* in broiler populations. Therefore, our genomic comparisons have been directed towards identification of any gene(s) that *S. agnetis* 908 could have acquired that facilitate the switch from involvement in cattle epithelial and mammary gland colonization and infection, to bone infections in chickens.

None of the gene islands (Table S2) or individual genes (Table S3) we identified as distinguishing isolate 908 from closely related cattle isolates is currently recognizable as a virulence marker, or that mediates tissue tropism. Previously we had identified 44 virulence genes in our annotation of the isolate 908 genome (26) and none of these genes is in the regions distinguishing the chicken and cattle isolates. The genomic analyses of the human-to-chicken jump for *S. aureus* was associated with acquisition of two prophage, two plasmids, and a pathogenicity island, the inactivation of several virulence determinants important to human pathogenesis, and enhanced resistance to chicken neutrophils (17). Thus, we expected to readily find genes, or gene clusters, in chicken isolates of *S. agnetis* associated with the jump to chickens from cattle. Most of the distinguishing gene islands in isolate 908 contain genes associated with mobile elements (prophage), but none are virulence determinants. We have unpublished evidence that isolate 908 is highly resistant to an immortalized chicken macrophage and are pursuing the genes for macrophage resistance. Since we have failed to identify unique virulence genes that distinguish the chicken isolate 908 from the cattle isolates we conclude that the basis for the jump from cattle to chickens is most likely the result of small alterations (i.e., missense or regulatory mutations) in a few virulence-associated factors. Hypervirulence of isolate 908 in chickens could be from a single amino acid change. Hypervirulence of isolates of *Campylobacter jejuni*, were demonstrated to result from a single substitution in an outer membrane protein, resulting in induction of spontaneous abortions in sheep (64). Therefore, further fine-level comparisons or directed genome evolution (64) will be required to dissect how this emerging pathogen has evolved and diversified from cattle mastitis to chicken bone pathogen.

## Materials and Methods

### Reference genomes

Isolate designations and host sources are provided in Table 1 and the details of the genome assemblies and accessions in NCBI are in Table S1. Chicken isolate 908 was from necrotic femoral lesions, while NDYM, NEDS and NEFX were from tissue samples from deceased broilers in Denmark. Isolates NDYM and NEDS were from the same broiler. The cattle isolate CBMRN was isolated from milk of a cow with subclinical mastitis that was enrolled in the Canadian Bovine Mastitis Research Network (CBMRN) cohort study (32). ATCC11249 represents the *S. hyicus* type strain isolated from a pig exudative epidermititis lesion (34).

### Bacterial strains

Genomes for ten *S. agnetis* isolates were newly assembled (Table S1), including nine cattle isolates, and one chicken isolate. Chicken isolate 1416 was isolated from a necrotic femoral lesion of a lame bird in a commercial broiler operation in Arkansas. The nine cattle isolates were from a collection at the University of Missouri. Two isolates were skin isolates, one isolate was obtained from a pre-partum mammary secretion, and six isolates were obtained from the milk of cows with subclinical mastitis. All cattle isolates had been previously identified as *S. agnetis* based on partial DNA sequence of either elongation factor Tu (*tuf*) or 3-dehydroquinate dehydratase (*aroD*) (30). All isolates were archived at -80 ^o^C in 20-40% glycerol, maintained on Tryptic Soy Agar slants, and grown in Tryptic Soy Broth (TSB; Difco, Becton, Dickinson and Company, Franklin Lakes, NJ).

### Genome sequencing and assembly

DNA isolation was based on the method described by Dyer and Iandolo (65). Isolates were grown to mid log phase in TSB (40 ml) at 37 ^o^C with shaking, pelleted, and resuspended in 2.5 ml 30 mM TrisCl, 3 mM EDTA, 50 mM NaCl, 50 mM glucose, pH 7.5. Lysostaphin (Sigma-Aldrich, St. Louis, MO) was added to 20 µg/ml, and incubated at 37 ^o^C for 40-60 min. SDS was added to 0.5%, then the lysate was treated with RNAseA (Sigma-Aldrich) at 20 ug/ul for 30 min at 37 ^o^C, then Pronase E (Sigma-Aldrich) at 20 ug/ul for 30 min at 37 ^o^C. The lysate was then extracted successively with 50:48:2 phenol:CHCl_3_:isoamyl alcohol, and 24:1 CHCl_3_:isoamyl alcohol. DNA was then collected by ethanol precipitation. DNA was quantified by Hoechst 33258 fluorometry in a GloMax®-Multi Jr. (Promega Corp., Madison, WI), and DNA integrity verified by agarose gel electrophoresis. Purified DNA from each isolate was submitted to the Research Technology Support Facility Genomics Core at Michigan State University for barcoded-library construction, pooled and subjected to 2×251 sequencing on an Illumina MiSeq. For draft genome assemblies the MiSeq reads were assembled using the *de novo* pipeline in Lasergene NGen ver. 13.0 (DNAStar, Madison, WI). For isolates 1379 and 1416 we produced finished genomes by hybrid assemblies of MiSeq and Oxford Nanopore MinION long reads. Long reads were either from barcoded or rapid kit libraries prepared and sequenced on Minion v9.3 flow cells (Oxford Nanopore Technologies, Oxford Science Park, UK) according to the manufacturer’s recommendations. Minion reads were filtered with a custom script to filter reads for length and average Q-score, prior to assembly. For isolate 1379, we filtered for length >=2000 bases and Qscore >=13. For isolate 1416, we filtered for length >=5000 bases and Qscore >16. Nanopore reads and MiSeq reads were assembled using the Unicycler ver. 0.4.6.0 pipeline on Galaxy (https://usegalaxy.org) using the Bold bridging mode. All assemblies and sequence reads have been deposited in NCBI and the accession and biosample identifiers are in Table S1.

### Genome annotation and phylogenetic comparison

The assembled genome sequences were compared with chicken isolate 908 using BLASTN implemented in CGViewer (37) to identify regions missing in one or more genome. Specific gene regions were annotated using either the BASys server at http://www.basys.ca (66) or the Rapid Annotation using System Technology (RAST) server at http://rast.nmpdr.org (67). Unique genes were verified by TBLASTN comparisons and reciprocal gene-by-gene BLASTP comparisons using the SEED server (68). Unique genes were further annotated using the KEGG (Kyoto Encyclopedia of Genes and Genomes) website at https://www.genome.jp/kegg (69). Prophage identification was performed using PHASTER (PHAge Search Tool Enhanced Release) at http://phaster.ca (70).

### Phylogenetic Analyses

For MLST analysis gene coding sequences were aligned and trimmed in MegAlign (DNAStar) then concatenated. Clustal Omega implemented in MegAlignPro (DNAStar) was used to generate phylogenetic trees. Consensus neighbor-joining trees with 2500 bootstrap replications were constructed based on the alignments. Genome-to-Genome Distance Calculator (GGDC) http://ggdc.dsmz.de/ggdc.php (35) was used to generate whole genome BLAST distance values. These distance calculations were used to generate a phylogenetic tree using the neighbor-joining method as implemented at http://trex.uqam.ca. Trees were rendered and rerooted in Archeopteryx 0.9901 (71).

## Acknowledgements

This research was partially supported by grants from the Arkansas Biosciences Institute, Chr Hansen, Zinpro LLC, and Cobb Vantress, Inc. The funders had no role in the design of this study, the interpretation of the results, or the contents of this manuscript.

