## Supplemental Tables 1 and 2 for "Whole genome comparisons of *Staphylococcus agnetis* isolates from cattle and chickens"

**Table S1.** Species, strain designation, genome assembly data, and accession number for isolates used in this study. Reference genomes were obtained from NCBI. Finished genomes utilized hybrid assemblies of Illumina and long reads (908- Pacific Biosciences; 1416 & 1379- Nanopore), while draft assemblies were *de novo* assemblies of Illumina or 454 short reads. Assembly data, where relevant, includes total count of Illumina short reads and filtered long reads (see details in Materials and Methods for filtering). For finished genomes where plasmids were identified, the contigs and lengths of the main chromosome and plasmids are listed separately. For the new draft genomes we include the approximate genome coverage in Illumina reads assuming a genome size of 2.5 Mbp, and the N50 of the final assembly. The NCBI Accession column contains Accession (A:) and Biosample (B:) for the genome data.

| **Species** | **Strain** | **Assembly Data** | | | | | | **NCBI Accession** | **Citation** |
| --- | --- | --- | --- | --- | --- | --- | --- | --- | --- |
|  |  | **Illumina Total Reads**  **(Coverage)** | **Filtered Long Reads**  **(size range)** | **Contigs** | **Length (Mbp)** | | |  |  |
| Reference genomes from NCBI | | | | | | | | | |
| *S. agnetis* | 908 | 8,476,030 | 66332 | 1 | 2.474 | | | CP009623.1 | (1) |
|  |  |  |  | 3 | 0.029, 0.003, 0.0022 | | |  |  |
| *S. agnetis* | CBMRN 20813338 |  |  | 45 | 2.416 | | | NZ_JPRT00000000.1 | (2) |
| *S. hyicus* | ATCC 11249 |  |  | 1 | 2.472 | | | NZ_CP008747.1 | (3) |
| *S. agnetis* | 722_260714_1_8_heart |  |  | 69 | 2.530 | | | NZ_NDYM00000000.1 | (4) |
| *S. agnetis* | 722_230714_2_5_spleen |  |  | 73 | 2.528 | | | NZ_NEDS00000000.1 |  |
| *S. agnetis* | 723_310714_2_2_spleen |  |  | 68 | 2.528 | | | NZ_NEFX00000000.1 |  |
| *S. agnetis* | 12B |  |  | 1 | 2.345 | | | GCA_007814015.1 | none |
| *S. agnetis* | DSM_23656 |  |  | 164 | 2.491 | | | GCA_002901865.1 | none |
| *S. agnetis* | SNUC_5151 |  |  | 31 | 2.387 | | | GCA_003040805.1 | (5) |
| *S. agnetis* | SNUC_4805 |  |  | 61 | 2.429 | | | GCA_003040835.1 | (5) |
| *S. agnetis* | SNUC_1371 |  |  | 54 | 2.407 | | | GCA_003040965.1 | (5) |
| *S. agnetis* | 3682 |  |  | 34 | 2.503 | | | GCA_007097565.1 | none |
| *S. agnetis* | 33 |  |  | 52 | 2.505 | | | GCA_002994165.1 | none |
| *S. agnetis* | 59 |  |  | 53 | 2.398 | | | GCA_002994125.1 | none |
| *S. agnetis* | SNUC_5631 |  |  | 69 | 2.342 | | | GCA_003040855.1 | (5) |
| *S. agnetis* | 43 |  |  | 58 | 2.450 | | | GCA_002994105.1 | none |
| *S. agnetis* | 59 |  |  | 58 | 2.476 | | | GCA_002994085.1 | none |
| *S. agnetis* | 6 |  |  | 58 | 2.476 | | | GCA_002994145.1 |  |
| *S. agnetis* | SNUC_3836 |  |  | 86 | 2.396 | | | GCA_003041195.1 | (5) |
| *S. agnetis* | SNUC_4051 |  |  | 95 | 2.488 | | | GCA_003040875.1 | (5) |
| *S. agnetis* | SNUC_719 |  |  | 101 | 2.401 | | | GCA_003041015.1 | (5) |
| *S. agnetis* | SNUC_2493 |  |  | 74 | 2.366 | | | GCA_003040935.1 | (5) |
| *S. agnetis* | SNUC_2265 |  |  | 101 | 2.453 | | | GCA_003041215.1 | (5) |
| *S. agnetis* | SNUC_725 |  |  | 111 | 2.398 | | | GCA_003040995.1 | (5) |
| *S. agnetis* | SNUC_3261 |  |  | 87 | 2.362 | | | GCA_003040915.1 | (5) |
| *S. agnetis* | SNUC_1383 |  |  | 104 | 2.435 | | | GCA_003040955.1 | (5) |
| *S. agnetis* | SNUC_3610 |  |  | 556 | 2.584 | | | GCA_003040895.1 | (5) |
| Finished genomes using long and short reads from this work | | | | | | | | | |
| *S. agnetis* | 1379 | 2,108,258  (210x) | 824  (2000-135765) | 1 | 2.449 | | | A: CP045927  B: SAMN13231958 |  |
| *S. agnetis* | 1416 | 3,167,394  (316x) | 53314 (5000-88691) | 1 | 2.448 | | | A: WMFQ00000000  B: SAMN13231956 |  |
|  |  |  |  | 2 | 0.059, 0.028 | | |  |  |
| Draft assemblies from short reads from this work | | | | | | **N50** | |  | |
| *S. agnetis* | 1383 | 1,659,666  (165x) |  | 43 | 2.381 | 111887 | | A: WMFO00000000  B: SAMN13231960 |  |
| *S. agnetis* | 1384 | 1,983,900  (198x) |  | 150 | 2.580 | 162149 | | A: WMFN00000000  B: SAMN13231961 |  |
| *S. agnetis* | 1385 | 2,170,488  (217x) |  | 328 | 2.581 | 55411 | | A: WMFM00000000  B: SAMN13231962 |  |
| *S. agnetis* | 1387 | 1,493,926  (149x) |  | 101 | 2.557 | | 84228 | A: WMFL00000000  B: SAMN13231963 |  |
| *S. agnetis* | 1389 | 1,594,062  (159x) |  | 127 | 2.547 | | 87399 | A: WMFK00000000  B: SAMN13231964 |  |
| *S. agnetis* | 1390 | 1,084,244  (108x) |  | 200 | 2.421 | | 47684 | A: WMFJ00000000  B: SAMN13231965 |  |
| *S. agnetis* | 1391 | 1,792,710  (179x) |  | 98 | 2.456 | | 51409 | A: WMFI00000000  B: SAMN13231969 |  |
| *S. agnetis* | 1392 | 1,569,576  (156x) |  | 62 | 2.443 | | 85109 | A: WMFH00000000  B: SAMN13231966 |  |

Table S2. BLASTP annotation and TBLASTN search of related *S. agnetis* genomes for predicted protein sequences from the five islands identified in *S. agnetis* 908. The predicted proteins from the RAST annotation (RAST peg) were annotated using BLASTP (eval 0.001) for Uniprot ID or NCBI ID, Protein name and the Organism, of the best match. TBLASTN results were for genomes of *S. agnetis* isolates: Danish (chicken draft genomes NEFX, NEDS, NDYM), 1416 (finished genome commercial broiler), 1379 (finished genome cattle isolate) and Bovine (31 bovine isolate genomes; the 9 draft genomes 1383, 1384, 1385, 1387, 1389, 1390, 1391, and 1392; plus the 22 listed under Reference genomes from NCBI in table S1 and Table 1). Numbers reported for TBLASTN were the eval of the best match (none is no significant match).

| RAST peg | Island | BLASTP | | | TBLASTN | | | |
| --- | --- | --- | --- | --- | --- | --- | --- | --- |
|  |  | ID | Protein names | Organism | Danish | 1416 | 1379 | Bovine |
| 167 | 1 | BLAI_STAAU | Penicillinase repressor (Beta-lactamase repressor protein) (Regulatory protein BlaI) | Staphylococcus aureus | none | none | none | 2E-71 |
| 168 | 1 | WP_001096367.1 | MULTISPECIES: beta-lactam sensor/signal transducer BlaR1 | Staphylococcus | none | none | none | 2E-99 |
| 169 | 1 | B1MCI3_MYCA9 | Beta-lactamase (EC 3.5.2.6) | Mycobacterium abscessus (strain ATCC 19977 / DSM 44196 / CIP 104536 / JCM 13569 / NCTC 13031 / TMC 1543) | none | none | none | 4E-139 |
| 170 | 1 | A0A0H2WWI3_STAAC | Uncharacterized protein | S. aureus (strain COL) | none | none | 1E-25 | 8E-63 |
| 171 | 1 | WP_002457078.1 | MULTISPECIES: hypothetical protein | Staphylococcus | none | none | none | none |
| 172 | 1 | WP_000410574.1 | MULTISPECIES: transposase | Staphylococcus | none | none | none | 6E-50 |
| 173 | 1 | XERC_MYCTU | Tyrosine recombinase XerC | Mycobacterium tuberculosis (strain ATCC 25618 / H37Rv) | none | none | 7E-09 | 0 |
| 174 | 1 | XERC_ECOLI | Tyrosine recombinase XerC | Escherichia coli (strain K12) | 1E-17 | none | 1E-17 | 0 |
| 175 | 1 | A0A0H2WWI3_STAAC | Uncharacterized protein | S. aureus (strain COL) | none | none | 2E-42 | 5E-43 |
| 176 | 1 | XERD_ECOLI | Tyrosine recombinase XerD | Escherichia coli (strain K12) | none | none | 0 | 0 |
| 177 | 1 | WP_047531257.1 | MULTISPECIES: hypothetical protein | Staphylococcus | none | none | none | 3E-78 |
| 178 | 1 | ALN76009.1 | putative membrane protein | Staphylococcus phage B166 | none | none | 2E-15 | 5E-21 |
| 179 | 1 | ARM67752.1 | hypothetical protein | Staphylococcus phage IME1318_01 | none | none | none | 4E-24 |
| 180 | 1 | WP_053028439.1 | MULTISPECIES: ImmA/IrrE family metallo-endopeptidase |  | none | none | none | 4E-05 |
| 181 | 1 | WP_002442276.1 | MULTISPECIES: XRE family transcriptional regulator | Staphylococcus | none | none | none | none |
| 182 | 1 | Y474_MYCTU | Uncharacterized HTH-type transcriptional regulator Rv0474 | M. tuberculosis (strain ATCC 25618 / H37Rv) | none | none | none | none |
| 183 | 1 | WP_062078533.1 | MULTISPECIES: antirepressor | Staphylococcus | none | none | 2E-30 | 6E-29 |
| 184 | 1 | WP_107362097.1 | MULTISPECIES: hypothetical protein | Staphylococcus | none | none | none | 1E-40 |
| 185 | 1 | WP_096608432.1 | MULTISPECIES: hypothetical protein | Staphylococcus | none | none | none | none |
| 186 | 1 | WP_060550713.1 | hypothetical protein | S. agnetis | none | none | 1E-36 | 2E-35 |
| 187 | 1 | WP_105996740.1 | hypothetical protein | Staphylococcus chromogenes | none | none | 6E-18 | 1E-21 |
| 188 | 1 | WP_060550714.1 | DUF1108 family protein |  | none | none | 2E-50 | 2E-49 |
| 189 | 1 | WP_105995253.1 | MULTISPECIES: DUF2483 domain-containing protein | Staphylococcus | none | none | 2E-40 | 2E-39 |
| 190 | 1 | WP_049395062.1 | MULTISPECIES: single-stranded DNA-binding protein | Staphylococcus | none | none | none | 6E-48 |
| 191 | 1 | SSB_HELPY | Single-stranded DNA-binding protein (SSB) | Helicobacter pylori (strain ATCC 700392 / 26695) (Campylobacter pylori) | none | none | 4E-18 | 1E-66 |
| 192 | 1 | WP_060550718.1 | hypothetical protein | S. agnetis | none | none | 1E-140 | 2E-134 |
| 193 | 1 | QSOX2_ARATH | Sulfhydryl oxidase 2 (EC 1.8.3.2) (Quiescin-sulfhydryl oxidase 2) (AtQSOX2) | Arabidopsis thaliana (Mouse-ear cress) | none | none | none | 9E-21 |
| 194 | 1 | WP_060550720.1 | DnaD domain protein | Staphylococcus | none | none | 6E-12 | 2E-137 |
| 195 | 1 | WP_060550721.1 | hypothetical protein | S. agnetis | none | none | 1E-25 | 3E-60 |
| 196 | 1 | Q9X4C9_GEOSE | Replicative DNA helicase (EC 3.6.4.12) | Geobacillus stearothermophilus (Bacillus stearothermophilus) | 2E-150 | none | 2E-151 | 0 |
| 197 | 1 | WP_060550723.1 | hypothetical protein | S. agnetis | none | none | 1E-18 | 5E-38 |
| 198 | 1 | WP_060550724.1 | MULTISPECIES: RusA family crossover junction endodeoxyribonuclease | Staphylococcus | none | none | 2E-46 | 1E-88 |
| 199 | 1 | WP_060550725.1 | MULTISPECIES: hypothetical protein | Staphylococcus | none | none | 2E-24 | 4E-77 |
| 200 | 1 | WP_060550726.1 | DUF3310 domain-containing protein | S. agnetis | none | none | none | 1E-82 |
| 201 | 1 | WP_060550727.1 | hypothetical protein | S. agnetis | none | none | none | 1E-81 |
| 202 | 1 | WP_060550728.1 | DUF3269 family protein | S. agnetis | none | none | 2E-39 | 2E-39 |
| 203 | 1 | WP_060550729.1 | hypothetical protein | S. agnetis | none | none | 3E-13 | 1E-22 |
| 204 | 1 | WP_060552463.1 | hypothetical protein | S. agnetis | none | none | 6E-33 | 6E-59 |
| 205 | 1 | WP_002504574.1 | MULTISPECIES: hypothetical protein | Staphylococcus | none | none | none | 7E-38 |
| 206 | 1 | WP_060550731.1 | hypothetical protein | S. agnetis | none | none | none | 2E-13 |
| 207 | 1 | WP_039645002.1 | MULTISPECIES: hypothetical protein | Staphylococcus | none | none | 3E-23 | 1E-22 |
| 208 | 1 | LCL3_YEAST | Probable endonuclease LCL3 (EC 3.1.-.-) (Long chronological lifespan protein 3) | Saccharomyces cerevisiae (strain ATCC 204508 / S288c) (Baker's yeast) | none | 3E-19 | 3E-72 | 2E-71 |
| 209 | 1 | DUT_MYCTU | Deoxyuridine 5'-triphosphate nucleotidohydrolase (dUTPase) (EC 3.6.1.23) (dUTP pyrophosphatase) | M. tuberculosis (strain ATCC 25618 / H37Rv) | none | none | none | 6E-78 |
| 210 | 1 |  | none |  | none | none | 1E-12 | 8E-17 |
| 211 | 1 | WP_085622443.1 | MULTISPECIES: hypothetical protein | Staphylococcus | none | none | 1E-33 | 3E-55 |
| 212 | 1 | WP_060550736.1 | hypothetical protein | S. agnetis | none | none | 7E-48 | 2E-49 |
| 213 | 1 | WP_060550737.1 | HNH endonuclease | S. agnetis | none | none | 2E-08 | 6E-07 |
| 214 | 1 | WP_060550738.1 | phage terminase small subunit P27 family |  | none | none | 4E-16 | 2E-16 |
| 215 | 1 | WP_046464342.1 | MULTISPECIES: terminase large subunit | Staphylococcus | none | none | 9E-51 | 6E-53 |
| 216 | 1 | INSI3_ECOLI | Transposase InsI for insertion sequence element IS30C | E. coli (strain K12) | none | none | none | 6E-34 |
| 217 | 1 | WP_014532495.1 | hypothetical protein | S. aureus | none | 4E-024 | none | none |
| 218 | 1 | O53337_MYCTU | Probable transposase | M. tuberculosis (strain ATCC 25618 / H37Rv) | none | 6E-37 | none | 1E-43 |
| 219 | 1 | WP_000166623.1 | MULTISPECIES: cystatin-like fold lipoprotein | Staphylococcus | none | none | none | 5E-27 |
| 220 | 1 | WP_000850646.1 | MULTISPECIES: hypothetical protein | Staphylococcus | none | none | none | 1E-61 |
| 221 | 1 | Q8DWM3_STRMU | Putative secreted antigen GbpB/SagA putative peptidoglycan hydrolase | Streptococcus mutans serotype c (strain ATCC 700610 / UA159) | 3E-20 | none | 2E-22 | 9E-136 |
| 222 | 1 | WP_000875438.1 | MULTISPECIES: hypothetical protein | Staphylococcus | none | none | none | 5E-161 |
| 223 | 1 | WP_000274938.1 | MULTISPECIES: hypothetical protein | Staphylococcus | none | none | none | none |
| 224 | 1 | FTSK_PSEAE | DNA translocase FtsK | Pseudomonas aeruginosa (strain ATCC 15692 / DSM 22644 / CIP 104116 / JCM 14847 / LMG 12228 / 1C / PRS 101 / PAO1) | 0 | none | 2E-09 | 0 |
| 225 | 1 | WP_031882373.1 | MULTISPECIES: hypothetical protein | Staphylococcus | none | none | none | none |
| 226 | 1 | WP_031882368.1 | MULTISPECIES: hypothetical protein | Staphylococcus | none | none | none | none |
| 227 | 1 | WP_060550739.1 | MULTISPECIES: hypothetical protein | Staphylococcus | none | none | none | none |
| 228 | 1 | YDDE_BACSU | Uncharacterized protein YddE | Bacillus subtilis (strain 168) | 0 | 0 | none | 0 |
| 229 | 1 | WP_000358151.1 | MULTISPECIES: hypothetical protein | Staphylococcus | none | none | none | 2E-42 |
| 230 | 1 | WP_000369240.1 | MULTISPECIES: hypothetical protein | Staphylococcus | none | none | none | 1E-20 |
| 231 | 1 | WP_060550741.1 | MULTISPECIES: conjugal transfer protein | Staphylococcus | none | none | none | 5E-123 |
| 232 | 1 | WP_001059817.1 | hypothetical protein | S. aureus | none | none | none | none |
| 233 | 1 | WP_050438939.1 | MULTISPECIES: replication initiation factor domain-containing protein | Staphylococcus | 3E-105 | none | none | 3E-100 |
| 234 | 1 | WP_001007543.1 | MULTISPECIES: hypothetical protein | Staphylococcus | none | none | none | 5E-09 |
| 235 | 1 | WP_000595049.1 | MULTISPECIES: DUF961 domain-containing protein | Staphylococcus | none | none | none | 2E-05 |
| 236 | 1 | WP_000180797.1 | MULTISPECIES: hypothetical protein | Staphylococcus | none | none | none | 2E-36 |
| 237 | 1 | WP_046464342.1 | MULTISPECIES: terminase large subunit | Staphylococcus | none | none | 7E-35 | 1E-34 |
| 238 | 1 | WP_047210745.1 | MULTISPECIES: hypothetical protein | Staphylococcus | none | none | none | none |
| 239 | 1 | WP_047210746.1 | MULTISPECIES: phage portal protein | Staphylococcus | none | none | 4E-47 | 1E-46 |
| 240 | 1 | PRO_BPHK7 | Prohead protease (GP4) | Enterobacteria phage HK97 (Bacteriophage HK97) | none | none | 3E-44 | 4E-43 |
| 241 | 1 | WP_060804357.1 | MULTISPECIES: phage major capsid protein | Staphylococcus | none | none | 7E-37 | 1E-37 |
| 242 | 1 | KKI61673.1 | hypothetical protein UF68_0775 | Staphylococcus warneri | none | none | 0.000003 | 2E-05 |
| 243 | 1 | WP_104681641.1 | MULTISPECIES: phage gp6-like head-tail connector protein | Staphylococcus | none | none | none | none |
| 244 | 1 | CRV14363.1 | Bacteriophage head-tail adaptor | Streptococcus equi subsp. equi | none | none | none | none |
| 245 | 1 | MOK13_SCHPO | Cell wall alpha-1,3-glucan synthase mok13 (EC 2.4.1.183) | Schizosaccharomyces pombe (strain 972 / ATCC 24843) (Fission yeast) | none | none | none | none |
| 246 | 1 | WP_060550749.1 | hypothetical protein | S. agnetis | none | none | none | none |
| 247 | 1 | WP_053017034.1 | MULTISPECIES: hypothetical protein | Staphylococcus | none | none | 3E-09 | 4E-11 |
| 248 | 1 | WP_060550751.1 | hypothetical protein | S. agnetis | none | none | none | none |
| 249 | 1 | YP_008853736.1 | hypothetical protein phiRS7_0013 | Staphylococcus phage phiRS7 | none | none | none | none |
| 250 | 1 | LYTM_STAA8 | Glycyl-glycine endopeptidase LytM (EC 3.4.24.75) (Autolysin LytM) | S. aureus (strain NCTC 8325) | none | 0 | 0 | 0 |
| 251 | 1 | REH96631.1 | phage tail protein | Staphylococcus felis | none | none | 3E-93 | 5E-94 |
| 252 | 1 | REI17843.1 | hypothetical protein DOS74_03015 | S. felis | 0 | 0 | 0 | 0 |
| 253 | 1 |  | none |  | none | none | none | none |
| 254 | 1 | WP_103210803.1 | CHAP domain-containing protein | S. felis | none | none | 0.0003 | 0.001 |
| 255 | 1 | PABPA_XENLA | Polyadenylate-binding protein 1-A (PABP-1-A) (Poly(A)-binding protein 1-A) (xPABP1-A) (Cytoplasmic poly(A)-binding protein 1-A) | Xenopus laevis (African clawed frog) | none | none | 1E-47 | 3E-48 |
| 256 | 1 | Q708K1_9CAUD | Holin | Streptococcus phage EJ-1 | none | none | none | 3E-90 |
| 257 | 1 | B8QIR1_9CAUD | LysH5 | Staphylococcus phage phiH5 | 1E-50 | none | 5E-52 | 0 |
| 258 | 1 | SOFIC_SHEON | Adenosine monophosphate-protein transferase SoFic (EC 2.7.7.n1) (AMPylator SoFic) | Shewanella oneidensis (strain MR-1) | none | none | none | 0 |
| 259 | 1 | A0A0H2UNK7_STRPN | Uncharacterized protein | Streptococcus pneumoniae serotype 4 (strain ATCC BAA-334 / TIGR4) | none | none | 8E-84 | 1E-83 |
| 260 | 1 | WP_060550761.1 | DUF2273 domain-containing protein | S. agnetis | none | none | 5E-28 | 1E-26 |
| 261 | 1 | WP_060550762.1 | alkaline shock response membrane anchor protein AmaP | S. agnetis | none | none | 1E-90 | 2E-91 |
| 974 | 2 | WP_060551312.1 | DUF2335 domain-containing protein | S. agnetis | none | none | none | none |
| 975 | 2 | WP_064782972.1 | MULTISPECIES: hypothetical protein | Staphylococcus | none | none | none | none |
| 976 | 2 | WP_060551314.1 | CPBP family intramembrane metalloprotease | S. agnetis | 2E-49 | none | 7E-08 | 2E-08 |
| 977 | 2 | LYTM_STAA8 | Glycyl-glycine endopeptidase LytM | S. aureus (strain NCTC 8325) | none | none | 1E-36 | 1E-58 |
| 978 | 2 | WP_096559817.1 | MULTISPECIES: PTS mannose transporter subunit IID | Staphylococcus | none | none | none | none |
| 979 | 2 | WP_115341903.1 | MULTISPECIES: DUF2951 family protein | Staphylococcus | none | none | none | none |
| 980 | 2 | WP_060551318.1 | hypothetical protein | S. agnetis | none | none | 0.00007 | none |
| 981 | 2 |  | none |  | none | none | none | none |
| 982 | 2 | WP_060551319.1 | hypothetical protein | S. agnetis | none | 0 | 0 | 0 |
| 983 | 2 | REH96631.1 | phage tail protein | S. felis | none | none | 2E-105 | 6E-106 |
| 984 | 2 | MEPM_ECOLI | Murein DD-endopeptidase MepM (EC 3.4.24.-) (Murein hydrolase MepM) (ORFU) | E. coli (strain K12) | 0 | none | 0 | 0 |
| 985 | 2 | WP_107371935.1 | MULTISPECIES: hypothetical protein | Staphylococcus | none | none | none | none |
| 986 | 2 | WP_096601407.1 | MULTISPECIES: hypothetical protein | Staphylococcus | none | none | none | none |
| 987 | 2 | WP_086375142.1 | MULTISPECIES: hypothetical protein | Staphylococcus | none | none | 0.0003 | 2E-06 |
| 988 | 2 | WP_060551323.1 | phage tail protein | S. agnetis | none | none | 4E-37 | 1E-36 |
| 989 | 2 | WP_060552487.1 | hypothetical protein | S. agnetis | none | none | none | 1E-08 |
| 990 | 2 | WP_070671866.1 | MULTISPECIES: hypothetical protein | Staphylococcus | none | none | none | 7E-30 |
| 991 | 2 | WP_107371940.1 | MULTISPECIES: hypothetical protein | Staphylococcus | none | none | none | none |
| 992 | 2 | WP_060551326.1 | MULTISPECIES: phage gp6-like head-tail connector protein | Staphylococcus | none | none | none | 7E-21 |
| 993 | 2 | WP_019168228.1 | MULTISPECIES: phage major capsid protein | Staphylococcus | none | none | none | 1E-80 |
| 994 | 2 | CLPP1_MYCTU | ATP-dependent Clp protease proteolytic subunit 1 (EC 3.4.21.92) (Endopeptidase Clp 1) | M. tuberculosis (strain ATCC 25618 / H37Rv) | none | none | 0.0008 | 2E-21 |
| 995 | 2 | WP_060551328.1 | phage portal protein | S. agnetis | none | none | none | 2E-102 |
| 996 | 2 | WP_060551329.1 | terminase large subunit | S. agnetis | none | none | 9E-35 | 6E-174 |
| 997 | 2 | WP_060551330.1 | MULTISPECIES: hypothetical protein | Staphylococcus | none | none | none | 4E-32 |
| 998 | 2 | ATD3C_HUMAN | ATPase family AAA domain-containing protein 3C | Homo sapiens (Human) | none | none | 1E-10 | 4E-21 |
| 999 | 2 | WP_060551331.1 | antitoxin HicB | S. agnetis | none | none | none | 9E-80 |
| 1000 | 2 | WP_037567940.1 | type II toxin-antitoxin system HicA family toxin | S. agnetis | none | none | none | 1E-33 |
| 1001 | 2 | WP_115056413.1 | transposase family protein | S. aureus | none | none | 1E-15 | 5E-53 |
| 1002 | 2 | WP_037572018.1 | MULTISPECIES: hypothetical protein | Staphylococcus | none | none | 4E-15 | 4E-14 |
| 1003 | 2 | WP_085622443.1 | MULTISPECIES: hypothetical protein | Staphylococcus | none | none | 2E-64 | 2E-63 |
| 1004 | 2 | YP_240224.1 | ORF204 | Staphylococcus virus EW | none | none | 3E-13 | 2E-09 |
| 1005 | 2 | WP_060551334.1 | hypothetical protein | S. agnetis | none | none | none | 1E-28 |
| 1006 | 2 | WP_060551335.1 | hypothetical protein | S. agnetis | none | none | 7E-34 | 2E-59 |
| 1007 | 2 | LCL3_YEAST | Probable endonuclease LCL3 | S. cerevisiae (strain ATCC 204508 / S288c) | 5E-09 | 1E-12 | 2E-65 | 2E-71 |
| 1008 | 2 | WP_060552490.1 | hypothetical protein | S. agnetis | none | none | 1E-30 | 8E-57 |
| 1009 | 2 | WP_060551337.1 | hypothetical protein | S. agnetis | none | none | 3E-23 | 2E-34 |
| 1010 | 2 | WP_060551338.1 | DUF3269 family protein | S. agnetis | none | none | 2E-39 | 2E-39 |
| 1011 | 2 | WP_060551339.1 | hypothetical protein | S. agnetis | none | none | none | 2E-80 |
| 1012 | 2 | WP_107381354.1 | MULTISPECIES: DUF3310 domain-containing protein | Staphylococcus | none | none | none | 3E-44 |
| 1013 | 2 | WP_060551341.1 | hypothetical protein | S. agnetis | none | none | 2E-23 | 8E-27 |
| 1014 | 2 | WP_060551342.1 | hypothetical protein | S. agnetis | none | none | 7E-33 | 7E-71 |
| 1015 | 2 | WP_060550724.1 | MULTISPECIES: RusA family crossover junction endodeoxyribonuclease | Staphylococcus | none | none | 9E-47 | 2E-86 |
| 1016 | 2 | WP_085622430.1 | MULTISPECIES: hypothetical protein | Staphylococcus | none | none | 4E-40 | 1E-39 |
| 1017 | 2 | DNAC_BACSU | Replicative DNA helicase (EC 3.6.4.12) | Bacillus subtilis (strain 168) | 6E-46 | 0 | 0 | 0 |
| 1018 | 2 | WP_107372956.1 | MULTISPECIES: hypothetical protein | Staphylococcus | none | none | 5E-51 | 4E-59 |
| 1019 | 2 | WP_060551346.1 | DnaD domain protein | S. agnetis | none | none | 2E-31 | 9E-139 |
| 1020 | 2 | WP_060551347.1 | hypothetical protein | S. agnetis | none | none | 1E-126 | 6E-128 |
| 1021 | 2 | SSB_HELPY | Single-stranded DNA-binding protein | H. pylori (strain ATCC 700392 / 26695) | none | none | 2E-18 | 2E-62 |
| 1022 | 2 | WP_049395062.1 | MULTISPECIES: single-stranded DNA-binding protein | Staphylococcus | none | none | none | 5E-48 |
| 1023 | 2 | Q8Y589_LISMO | Lmo2180 protein | Listeria monocytogenes serovar 1/2a (strain ATCC BAA-679 / EGD-e) | none | none | none | 2E-14 |
| 1024 | 2 | WP_060551351.1 | DUF1108 family protein | S. agnetis | none | none | 3E-45 | 2E-50 |
| 1025 | 2 | WP_105996740.1 | hypothetical protein | S. chromogenes | none | none | 1E-09 | 3E-12 |
| 1026 | 2 | WP_060551352.1 | DUF771 domain-containing protein | S. agnetis | none | none | none | 1E-59 |
| 1027 | 2 | WP_060552492.1 | hypothetical protein | S. agnetis | none | none | none | 6E-16 |
| 1028 | 2 | WP_070626520.1 | MULTISPECIES: hypothetical protein | Staphylococcus | none | none | none | none |
| 1029 | 2 | WP_070626519.1 | MULTISPECIES: hypothetical protein | Staphylococcus | none | none | none | 4E-20 |
| 1030 | 2 | WP_100008585.1 | DUF2829 domain-containing protein | Staphylococcus pseudintermedius | none | none | none | 1E-10 |
| 1031 | 2 | WP_070848627.1 | MULTISPECIES: hypothetical protein | Staphylococcus | none | none | none | none |
| 1032 | 2 | WP_107378107.1 | phage antirepressor Ant | S. chromogenes | none | none | 4E-12 | 1E-10 |
| 1033 | 2 | CGNL1_MOUSE | Cingulin-like protein 1 (Junction-associated coiled-coil protein) | Mus musculus (Mouse) | none | none | none | 2E-20 |
| 1034 | 2 | WP_047432554.1 | MULTISPECIES: XRE family transcriptional regulator | Staphylococcus | none | none | 2E-52 | 4E-51 |
| 1035 | 2 | Q5M4N8_STRT2 | Transcriptional regulator | Streptococcus thermophilus (strain ATCC BAA-250 / LMG 18311) | none | none | 1E-81 | 3E-80 |
| 1036 | 2 | WP_107365676.1 | MULTISPECIES: hypothetical protein | Staphylococcus | none | none | 3E-17 | 6E-17 |
| 1037 | 2 | WP_070848523.1 | MULTISPECIES: hypothetical protein | Staphylococcus | none | none | none | 3E-08 |
| 1038 | 2 | XERC_MYCTU | Tyrosine recombinase XerC | M. tuberculosis (strain ATCC 25618 / H37Rv) | 0.0001 | none | 6E-79 | 0 |
| 1179 | 3 | WP_032604072.1 | MULTISPECIES: hypothetical protein | Staphylococcus | none | none | none | none |
| 1180 | 3 | O06608_MYCTU | Probable PhiRv1 phage protein | M. tuberculosis (strain ATCC 25618 / H37Rv) | none | none | none | 1E-88 |
| 1181 | 3 | WP_107388288.1 | mobile element-associated protein | Staphylococcus chromogenes | none | none | 2E-168 | 1E-177 |
| 1182 | 3 | WP_070724664.1 | MULTISPECIES: DUF1474 family protein | Staphylococcus | none | none | 3E-17 | 8E-51 |
| 1183 | 3 | PWZ93695.1 | pathogenicity island protein | S. pseudintermedius | none | none | none | 6E-83 |
| 1184 | 3 | WP_015728770.1 | MULTISPECIES: hypothetical protein | Staphylococcus | none | none | 3E-36 | 6E-37 |
| 1185 | 3 | WP_060551480.1 | helix-turn-helix domain-containing protein | S. agnetis | none | none | none | 2E-56 |
| 1186 | 3 | WP_110168783.1 | phage repressor protein | S. pseudintermedius | none | none | 0.000007 | 4E-05 |
| 1187 | 3 | SUTR_ECOLI | HTH-type transcriptional regulator SutR (Sulfur utilization regulator) | E. coli (strain K12) | none | none | 0.000002 | 4E-07 |
| 1188 | 3 | WP_050439619.1 | MULTISPECIES: XRE family transcriptional regulator | Staphylococcus | none | none | none | none |
| 1189 | 3 | INTQ_ECOLI | Putative defective protein IntQ (Putative lambdoid prophage Qin defective integrase) | E. coli (strain K12) | 7E-15 | 3E-08 | 1E-10 | 0 |
| 1813 | 4 | O53337_MYCTU | Probable transposase | M. tuberculosis (strain ATCC 25618 / H37Rv) | none | 2E-09 | none | 5E-80 |
| 1814 | 4 | O53337_MYCTU | Probable transposase | M. tuberculosis (strain ATCC 25618 / H37Rv) | none | 1E-51 | none | 4E-68 |
| 1815 | 4 | WP_060552013.1 | MULTISPECIES: cystatin-like fold lipoprotein | Staphylococcus | none | none | none | 5E-28 |
| 1816 | 4 | WP_060552014.1 | hypothetical protein | S. agnetis | none | none | none | 4E-72 |
| 1817 | 4 | Q8DWM3_STRMU | Putative secreted antigen GbpB/SagA putative peptidoglycan hydrolase | S. mutans serotype c (strain ATCC 700610 / UA159) | none | none | 3E-21 | 2E-148 |
| 1818 | 4 | MAP1A_RAT | Microtubule-associated protein 1A (MAP-1A) [Cleaved into: MAP1A heavy chain; MAP1 light chain LC2] | Rattus norvegicus (Rat) | 0 | none | none | 0 |
| 1819 | 4 | WP_031838453.1 | MULTISPECIES: hypothetical protein | Staphylococcus | none | none | none | none |
| 1820 | 4 | WP_031838454.1 | MULTISPECIES: hypothetical protein | Staphylococcus | none | none | none | none |
| 1821 | 4 | FTSK_ECOLI | DNA translocase FtsK | E. coli (strain K12) | none | none | 2E-16 | 0 |
| 1822 | 4 | WP_031796727.1 | MULTISPECIES: hypothetical protein | Staphylococcus | none | none | none | none |
| 1823 | 4 | CXU05770.1 | transposon-related protein | S. aureus | none | none | none | 2E-06 |
| 1824 | 4 | O53463_MYCTU | Transcriptional regulatory protein | M. tuberculosis (strain ATCC 25618 / H37Rv) | 0 | none | none | 4E-102 |
| 1825 | 4 | WP_031796729.1 | MULTISPECIES: hypothetical protein | Staphylococcus | none | none | none | 3E-06 |
| 1826 | 4 | YDDE_BACSU | Uncharacterized protein YddE | B. subtilis (strain 168) | 0 | 0 | none | 0 |
| 1827 | 4 | WP_060552019.1 | MULTISPECIES: conjugal transfer protein | Staphylococcus | none | none | none | 6E-37 |
| 1828 | 4 | WP_060552020.1 | MULTISPECIES: hypothetical protein | Staphylococcus | none | none | none | 1E-27 |
| 1829 | 4 | WP_060552021.1 | MULTISPECIES: conjugal transfer protein | Staphylococcus | none | none | none | 2E-119 |
| 1830 | 4 | WP_060552022.1 | MULTISPECIES: replication initiation factor domain-containing protein | Staphylococcus | none | none | none | 0 |
| 1831 | 4 | WP_060552023.1 | MULTISPECIES: hypothetical protein | Staphylococcus | none | none | none | 5E-42 |
| 1832 | 4 | WP_060552024.1 | MULTISPECIES: DUF961 domain-containing protein | Staphylococcus | none | none | none | 6E-47 |
| 1833 | 4 | WP_060552025.1 | MULTISPECIES: hypothetical protein | Staphylococcus | none | none | none | 6E-33 |
| 1978 | 5 | WP_107387598.1 | MULTISPECIES: bacteriochlorophyll 4-vinyl reductase | Staphylococcus | none | none | none | 3E-21 |
| 1979 | 5 | WP_060552141.1 | type I restriction endonuclease subunit R | S. agnetis | 0 | none | none | 0 |
| 1980 | 5 | WP_082624774.1 | restriction endonuclease subunit S | S. agnetis | none | none | none | 1E-121 |
| 1981 | 5 | O33298_MYCTU | Possible type I restriction/modification system DNA methylase HsdM (M protein) (DNA methyltransferase) | M. tuberculosis (strain ATCC 25618 / H37Rv) | none | none | none | 0 |
| 1982 | 5 | WP_039643461.1 | MULTISPECIES: hypothetical protein | Staphylococcus | none | none | none | 4E-120 |
| 1983 | 5 | WP_107397199.1 | MULTISPECIES: ATP-binding protein | Staphylococcus | none | none | none | 3E-04 |

Table S3. SEED analysis of predicted polypeptides present in genomes of chicken isolates of *S. agnetis* but not in two representative cattle isolates. The SEED server was used to identify polypeptides present in the genomes of isolates 908 and 1416 but less than 50% conserved in isolates 1379 and CBMRN. Those polypeptides are identified as 908 source (2.4 Mbp chromosome, 29 kbp plasmid, or 2.2 kbp plasmid) RAST Gene ID, Start (bp), location in islands (Table S2 and text), predicted Polypeptide length, functional annotation from RAST and PGAP, and Percent identity in the other three genomes.

| Source | Gene ID | Start | Island | Polypeptide Length | RAST function | PGAP function | Percent identity | | |
| --- | --- | --- | --- | --- | --- | --- | --- | --- | --- |
|  |  |  |  |  |  |  | 1416 | 1379 | CBMRN |
| Hypothetical protein, unknown function | | | | | | | | | |
| 2.4 Mbp | 193 | 184575 | 1 | 68 | hypothetical protein | hypothetical protein | 100.0 | 0.0 | 0.0 |
| 2.4 Mbp | 199 | 187812 | 1 | 141 | hypothetical protein | hypothetical protein | 100.0 | 39.7 | 43.3 |
| 2.4 Mbp | 200 | 188231 | 1 | 137 | hypothetical protein | DUF3310 domain-containing protein | 100.0 | 0.0 | 0.0 |
| 2.4 Mbp | 206 | 190382 | 1 | 53 | hypothetical protein | hypothetical protein | 100.0 | 0.0 | 0.0 |
| 2.4 Mbp | 217 | 195730 | 1 | 47 | hypothetical protein | none | 100.0 | 0.0 | 0.0 |
| 2.4 Mbp | 220 | 196934 | 1 | 198 | hypothetical protein | hypothetical protein | 100.0 | 0.0 | 0.0 |
| 2.4 Mbp | 222 | 198551 | 1 | 644 | hypothetical protein | hypothetical protein | 100.0 | 0.0 | 0.0 |
| 2.4 Mbp | 223 | 200582 | 1 | 134 | hypothetical protein | hypothetical protein | 100.0 | 0.0 | 0.0 |
| 2.4 Mbp | 225 | 202351 | 1 | 111 | hypothetical protein | hypothetical protein | 100.0 | 0.0 | 0.0 |
| 2.4 Mbp | 226 | 202680 | 1 | 77 | hypothetical protein | hypothetical protein | 100.0 | 0.0 | 0.0 |
| 2.4 Mbp | 227 | 202914 | 1 | 73 | hypothetical protein | hypothetical protein | 100.0 | 0.0 | 0.0 |
| 2.4 Mbp | 229 | 205674 | 1 | 130 | hypothetical protein transposon-related | hypothetical protein | 100.0 | 0.0 | 0.0 |
| 2.4 Mbp | 230 | 206069 | 1 | 87 | hypothetical protein | hypothetical protein | 100.0 | 0.0 | 0.0 |
| 2.4 Mbp | 232 | 207485 | 1 | 40 | hypothetical protein | none | 100.0 | 0.0 | 0.0 |
| 2.4 Mbp | 234 | 208943 | 1 | 142 | hypothetical protein | hypothetical protein | 100.0 | 0.0 | 0.0 |
| 2.4 Mbp | 235 | 209390 | 1 | 108 | hypothetical protein | DUF961 domain-containing protein | 100.0 | 0.0 | 0.0 |
| 2.4 Mbp | 236 | 209868 | 1 | 95 | hypothetical protein transposon-related | hypothetical protein | 100.0 | 0.0 | 0.0 |
| 2.4 Mbp | 238 | 210856 | 1 | 62 | hypothetical protein | hypothetical protein | 100.0 | 0.0 | 0.0 |
| 2.4 Mbp | 242 | 214206 | 1 | 58 | hypothetical protein | none | 100.0 | 48.2 | 48.2 |
| 2.4 Mbp | 245 | 215001 | 1 | 142 | hypothetical protein | hypothetical protein | 100.0 | 0.0 | 0.0 |
| 2.4 Mbp | 246 | 215416 | 1 | 138 | hypothetical protein | hypothetical protein | 100.0 | 0.0 | 0.0 |
| 2.4 Mbp | 247 | 215836 | 1 | 286 | hypothetical protein | hypothetical protein | 100.0 | 28.1 | 28.8 |
| 2.4 Mbp | 248 | 216760 | 1 | 180 | hypothetical protein | hypothetical protein | 100.0 | 0.0 | 0.0 |
| 2.4 Mbp | 249 | 217314 | 1 | 56 | hypothetical protein | none | 100.0 | 0.0 | 0.0 |
| 2.4 Mbp | 250 | 217500 | 1 | 1935 | hypothetical protein | hypothetical protein | 100.0 | 47.2 | 47.0 |
| 2.4 Mbp | 251 | 223305 | 1 | 497 | hypothetical protein | hypothetical protein | 100.0 | 36.8 | 44.8 |
| 2.4 Mbp | 252 | 224797 | 1 | 1161 | hypothetical protein | hypothetical protein | 100.0 | 45.0 | 47.3 |
| 2.4 Mbp | 982 | 983724 | 2 | 1161 | hypothetical protein | hypothetical protein | 99.0 | 45.3 | 47.4 |
| 2.4 Mbp | 1023 | 1012617 | 2 | 162 | hypothetical protein | hypothetical protein | 100.0 | 0.0 | 32.7 |
| 2.4 Mbp | 1027 | 1013955 | 2 | 94 | hypothetical protein | hypothetical protein | 100.0 | 0.0 | 0.0 |
| 2.4 Mbp | 1028 | 1014277 | 2 | 81 | hypothetical protein | hypothetical protein | 100.0 | 0.0 | 0.0 |
| 2.4 Mbp | 1029 | 1014568 | 2 | 112 | hypothetical protein | hypothetical protein | 100.0 | 0.0 | 0.0 |
| 2.4 Mbp | 1032 | 1015489 | 2 | 196 | hypothetical protein | hypothetical protein | 100.0 | 25.3 | 0.0 |
| 2.4 Mbp | 1033 | 1016079 | 2 | 308 | hypothetical protein | hypothetical protein | 100.0 | 0.0 | 32.0 |
| 2.4 Mbp | 1573 | 1570963 |  | 41 | hypothetical protein | none | 100.0 | 0.0 | 0.0 |
| 2.2 kbp | 2444 | 346 |  | 209 | hypothetical protein | none | 100.0 | 0.0 | 0.0 |
| 2.2 kbp | 2445 | 1369 |  | 297 | hypothetical protein | none | 100.0 | 0.0 | 0.0 |
| 29 kbp | 2451 | 465 |  | 190 | hypothetical protein | hypothetical protein | 100.0 | 0.0 | 0.0 |
| 29 kbp | 2452 | 1095 |  | 56 | hypothetical protein | none | 100.0 | 0.0 | 0.0 |
| 29 kbp | 2454 | 3093 |  | 156 | hypothetical protein | hypothetical protein | 100.0 | 0.0 | 0.0 |
| 29 kbp | 2455 | 4136 |  | 209 | hypothetical protein | hypothetical protein | 100.0 | 0.0 | 0.0 |
| 29 kbp | 2456 | 5133 |  | 167 | hypothetical protein | hypothetical protein | 100.0 | 0.0 | 0.0 |
| 29 kbp | 2457 | 5783 |  | 90 | hypothetical protein | hypothetical protein | 100.0 | 0.0 | 0.0 |
| 29 kbp | 2458 | 6204 |  | 40 | hypothetical protein | none | 100.0 | 0.0 | 0.0 |
| 29 kbp | 2463 | 8646 |  | 64 | hypothetical protein | hypothetical protein | 100.0 | 0.0 | 0.0 |
| 29 kbp | 2467 | 10691 |  | 68 | hypothetical protein | none | 100.0 | 0.0 | 0.0 |
| 29 kbp | 2473 | 15056 |  | 171 | hypothetical protein | hypothetical protein | 100.0 | 26.1 | 27.6 |
| 29 kbp | 2475 | 16192 |  | 99 | hypothetical protein | hypothetical protein | 100.0 | 0.0 | 0.0 |
| 29 kbp | 2479 | 19409 |  | 102 | hypothetical protein | hypothetical protein | 100.0 | 0.0 | 0.0 |
| 29 kbp | 2482 | 24434 |  | 107 | hypothetical protein | hypothetical protein | 100.0 | 0.0 | 0.0 |
| 29 kbp | 2483 | 24800 |  | 72 | hypothetical protein | hypothetical protein | 100.0 | 0.0 | 0.0 |
| 29 kbp | 2484 | 25842 |  | 73 | hypothetical protein | hypothetical protein | 100.0 | 0.0 | 0.0 |
| 29 kbp | 2485 | 26220 |  | 170 | hypothetical protein | hypothetical protein | 100.0 | 0.0 | 0.0 |
| Phage function | | | | | | | | | |
| 2.4 Mbp | 190 | 182832 | 1 | 221 | hypothetical protein | single-stranded DNA-binding protein | 99.1 | 0.0 | 0.0 |
| 2.4 Mbp | 195 | 185614 | 1 | 114 | Phage protein | hypothetical protein | 100.0 | 42.2 | 45.2 |
| 2.4 Mbp | 197 | 187183 | 1 | 73 | Phage protein | hypothetical protein | 100.0 | 45.8 | 45.8 |
| 2.4 Mbp | 201 | 188638 | 1 | 165 | Phage protein | hypothetical protein | 100.0 | 0.0 | 0.0 |
| 2.4 Mbp | 205 | 190066 | 1 | 99 | Phage protein | hypothetical protein | 100.0 | 0.0 | 0.0 |
| 2.4 Mbp | 214 | 193559 | 1 | 157 | Phage terminase, small subunit | phage terminase small subunit P27 family | 100.0 | 32.4 | 31.9 |
| 2.4 Mbp | 215 | 194022 | 1 | 378 | Phage terminase, large subunit | hypothetical protein | 100.0 | 34.6 | 35.5 |
| 2.4 Mbp | 237 | 210239 | 1 | 202 | Phage terminase, large subunit | hypothetical protein | 100.0 | 37.3 | 37.3 |
| 2.4 Mbp | 239 | 211051 | 1 | 410 | Phage portal protein | phage portal protein | 100.0 | 31.8 | 32.1 |
| 2.4 Mbp | 240 | 212267 | 1 | 188 | hypothetical protein | HK97 family phage prohead protease | 100.0 | 29.9 | 49.7 |
| 2.4 Mbp | 241 | 212870 | 1 | 439 | hypothetical protein | phage major capsid protein | 100.0 | 28.1 | 28.1 |
| 2.4 Mbp | 243 | 214388 | 1 | 104 | hypothetical protein | phage gp6-like head-tail connector protein | 100.0 | 0.0 | 0.0 |
| 2.4 Mbp | 244 | 214681 | 1 | 108 | hypothetical protein | head-tail adaptor protein | 100.0 | 0.0 | 0.0 |
| 2.4 Mbp | 1011 | 1006618 | 2 | 164 | Phage protein | hypothetical protein | 96.9 | 0.0 | 0.0 |
| 2.4 Mbp | 1022 | 1011954 | 2 | 221 | hypothetical protein | single-stranded DNA-binding protein | 100.0 | 0.0 | 0.0 |
| 2.4 Mbp | 1030 | 1014889 | 2 | 78 | Phage protein | hypothetical protein | 100.0 | 0.0 | 0.0 |
| 2.4 Mbp | 1031 | 1015164 | 2 | 105 | Phage protein | hypothetical protein | 100.0 | 0.0 | 0.0 |
| Mobile elements, plasmid replication, or toxin/antitoxin | | | | | | | | | |
| 2.4 Mbp | 194 | 184817 | 1 | 267 | hypothetical protein | DnaD domain protein | 100.0 | 38.3 | 39.0 |
| 2.4 Mbp | 213 | 193034 | 1 | 131 | hypothetical protein | HNH endonuclease | 100.0 | 38.8 | 38.8 |
| 2.4 Mbp | 216 | 195429 | 1 | 100 | Mobile element protein | hypothetical protein | 100.0 | 0.0 | 0.0 |
| 2.4 Mbp | 218 | 195883 | 1 | 177 | hypothetical protein | transposase | 100.0 | 0.0 | 0.0 |
| 2.4 Mbp | 231 | 206334 | 1 | 359 | hypothetical protein transposon-related | conjugal transfer protein | 100.0 | 0.0 | 0.0 |
| 2.4 Mbp | 233 | 207638 | 1 | 380 | hypothetical protein | replication initiation factor domain-containing protein | 100.0 | 0.0 | 0.0 |
| p3.0 | 2449 | 1650 |  | 282 | Replication protein | none | 90.4 | 0.0 | 0.0 |
| 29 kbp | 2453 | 2344 |  | 187 | Replication protein | replication protein | 91.5 | 0.0 | 0.0 |
| 29 kbp | 2459 | 6633 |  | 86 | YefM protein (antitoxin to YoeB) | type II toxin-antitoxin system Phd/YefM family antitoxin | 100.0 | 0.0 | 0.0 |
| 29 kbp | 2460 | 6890 |  | 89 | mRNA interferase RelE | Txe/YoeB family addiction module toxin | 100.0 | 0.0 | 0.0 |
| 29 kbp | 2461 | 7587 |  | 185 | Mobile element protein | recombinase family protein | 100.0 | 0.0 | 0.0 |
| 29 kbp | 2465 | 10094 |  | 113 | hypothetical protein | type II toxin-antitoxin system PemK/MazF family toxin | 100.0 | 0.0 | 0.0 |
| 29 kbp | 2477 | 17332 |  | 41 | hypothetical protein | IS5/IS1182 family transposase | 100.0 | 0.0 | 0.0 |
| 29 kbp | 2480 | 19720 |  | 264 | Chromosome (plasmid) partitioning protein ParA | ParA family protein | 100.0 | 0.0 | 0.0 |
| 29 kbp | 2481 | 20999 |  | 326 | Replication initiation protein A | hypothetical protein | 100.0 | 34.9 | 34.9 |
| Other | | | | | | | | | |
| 2.4 Mbp | 209 | 191107 | 1 | 173 | Deoxyuridine 5'-triphosphate nucleotidohydrolase (EC 3.6.1.23), phage associated | dUTP pyrophosphatase | 100.0 | 0.0 | 0.0 |
| 2.4 Mbp | 219 | 196525 | 1 | 121 | hypothetical protein | cystatin-like fold lipoprotein | 100.0 | 0.0 | 0.0 |
| 2.4 Mbp | 221 | 197533 | 1 | 343 | hypothetical protein | mannosyl-glycoprotein endo-beta-N-acetylglucosamidase | 100.0 | 48.5 | 45.1 |
| 2.4 Mbp | 224 | 200988 | 1 | 453 | hypothetical protein | cell division protein FtsK | 100.0 | 26.9 | 26.9 |
| 2.4 Mbp | 228 | 203144 | 1 | 832 | hypothetical protein | AAA family ATPase | 100.0 | 0.0 | 0.0 |
| 2.4 Mbp | 1037 | 1018987 | 2 | 239 | Hypothetical SAR0365 homolog in superantigen-encoding pathogenicity islands SaPI | hypothetical protein | 100.0 | 0.0 | 0.0 |
| 29 kbp | 2462 | 8152 |  | 83 | hypothetical protein | XRE family transcriptional regulator | 100.0 | 0.0 | 0.0 |
| 29 kbp | 2464 | 9236 |  | 228 | hypothetical protein | CPBP family intramembrane metalloprotease | 100.0 | 31.4 | 31.4 |
| 29 kbp | 2466 | 10429 |  | 90 | Ribosyl nicotinamide transporter, PnuC-like | hypothetical protein | 100.0 | 0.0 | 0.0 |
| 29 kbp | 2469 | 11845 |  | 175 | Aspartate aminotransferase (EC 2.6.1.1) | N-acetyl-L,L-diaminopimelate aminotransferase | 100.0 | 45.2 | 45.2 |
| 29 kbp | 2470 | 12387 |  | 206 | N-acetyl-L,L-diaminopimelate aminotransferase (EC 2.6.1.-) | none | 100.0 | 41.5 | 41.3 |
| 29 kbp | 2471 | 13308 |  | 244 | hypothetical protein | CPBP family intramembrane metalloprotease | 100.0 | 27.3 | 31.1 |
| 29 kbp | 2472 | 14152 |  | 270 | hypothetical protein | lysophospholipase | 100.0 | 24.3 | 24.3 |
| 29 kbp | 2474 | 15617 |  | 191 | hypothetical protein | CPBP family intramembrane metalloprotease | 100.0 | 0.0 | 0.0 |
